# CRISPR knockout genome-wide screens identify the HELQ-RAD52 axis in regulating the repair of cisplatin-induced single stranded DNA gaps

**DOI:** 10.1101/2024.04.17.589988

**Authors:** Lindsey M. Pale, Jude B. Khatib, Claudia M. Nicolae, George-Lucian Moldovan

**Author notes:** Correspondence:* George-Lucian Moldovan.

## Abstract

Treatment with genotoxic agents, such as platinum compounds, is still the mainstay therapeutical approach for the majority of cancers. Our understanding of the mechanisms of action of these drugs is however imperfect, and continuously evolving. Recent advances in the field highlighted single stranded DNA (ssDNA) gap accumulation as a potential determinant underlying cisplatin chemosensitivity, at least in some genetic backgrounds, such as BRCA mutations. Cisplatin-induced ssDNA gaps form upon the arrest of replication forks at sites of cisplatin adducts, and restart of DNA synthesis downstream of the lesion through repriming catalyzed by the PRIMPOL enzyme. Here, we show that PRIMPOL overexpression in otherwise wildtype cells results in accumulation of cisplatin-induced ssDNA gaps without sensitizing cells to cisplatin, suggesting that ssDNA gap accumulation does not confer cisplatin sensitivity in BRCA-proficient cells. To understand how ssDNA gaps may cause cellular sensitivity, we employed CRISPR-mediated genome-wide genetic screening to identify factors which enable the cytotoxicity of cisplatin-induced ssDNA gaps. We found that the helicase HELQ specifically suppresses cisplatin sensitivity in PRIMPOL-overexpressing cells, and this is associated with reduced ssDNA accumulation. We moreover identify RAD52 as a mediator of this pathway, and show that RAD52 promotes ssDNA gap accumulation through a BRCA-mediated mechanism. Our work identified the HELQ-RAD52-BRCA axis as a regulator of ssDNA gap processing, shedding light on the mechanisms of cisplatin sensitization in cancer therapy.

## Introduction

Despite advances in targeted therapies, a significant proportion of cancers are still treated with conventional chemotherapy aimed at inducing DNA damage in cancer cells. A classic example is represented by cisplatin and derivative platinum compounds, widely used in cancer treatment (Zhang et al., 2022). Cisplatin creates intrastrand DNA crosslinks and other types of DNA lesions, arresting replication forks and thus interfering with DNA synthesis in rapidly proliferating cancer cells.

Upon fork arrest at these lesions, several different processes can occur. Arrested forks can reverse, by annealing of the nascent strand of the sister chromatids, catalyzed by DNA translocases including SMARCAL1, ZRANB3 and HLTF (Bhat and Cortez, 2018; Quinet et al., 2017). This process stabilizes the fork structure, allowing time for lesion excision and also providing an opportunity for fork restart using the nascent strand of the sister chromatid as a temporary template. The resolution of the reversed fork structure, catalyzed by various helicases, allows resumption of replication from the original template. Since it exposes a double stranded end, the reversed fork structure needs to be protected against nucleolytic degradation. The BRCA tumor suppressor pathway is essential for this, by catalyzing RAD51 filament formation on the reversed arm. In BRCA-deficient cells, MRE11, EXO1 and other nucleases degrade the nascent DNA starting on the reversed arm, resulting in genomic instability (Bhat and Cortez, 2018; Dhoonmoon et al., 2022; Guillemette et al., 2015; Kolinjivadi et al., 2017; Lemacon et al., 2017; Mijic et al., 2017; Quinet et al., 2017; Ray Chaudhuri et al., 2016; Schlacher et al., 2011; Taglialatela et al., 2017; Thakar and Moldovan, 2021).

Alternatively, the stalled fork can be restarted through the activity of the primase-polymerase PRIMPOL, which is recruited downstream of the arresting lesion. As the name implies, PRIMPOL is able to start replication without a need for an existing primer, but instead it is able to synthesize the primer itself (Bianchi et al., 2013; Mouron et al., 2013). Subsequently, PRIMPOL is exchanged with a replicative polymerase to resume normal DNA synthesis. This process leaves behind a single stranded DNA (ssDNA) gap, which needs to be filled at a later time. The BRCA pathway was shown to be important for gap suppression (Cong et al., 2021a; Hale et al., 2023; Panzarino et al., 2021; Quinet et al., 2021; Quinet et al., 2020; Tirman et al., 2021). This potentially reflects a role for BRCA-mediated recombination using the nascent strand of the sister chromatid as template for gap filling. It is also possible that the BRCA pathway suppresses the engagement of nucleases on ssDNA structures, similar to its activity in protecting reversed forks against nucleolytic degradation. In line with this, it was recently shown that nucleases including MRE11 and EXO1 are expanding ssDNA gaps in BRCA-deficient cells (Hale et al., 2023; Tirman et al., 2021). Alternatively, ssDNA gaps can also be filled through translesion DNA synthesis (TLS), a process which involves specialized, low-fidelity polymerases able to bypass DNA lesions and extend beyond the lesion on the undamaged template. BRCA-deficient cells are reliant on TLS for gap filling (Taglialatela et al., 2021; Tirman et al., 2021).

Failure to stabilize arrested forks or endonucleolytic processing of the ssDNA region at the arresting site results in formation of double stranded DNA breaks (DSBs) (Zeman and Cimprich, 2014), which are highly cytotoxic structures. Replication-associated DSBs can however be repaired through BRCA-mediated recombination. In BRCA-deficient cells, more mutagenic end-joining mechanisms are employed for DSB repair (Shrivastav et al., 2008).

The BRCA pathway is frequently inactivated in tumors, through both germline mutations (in individuals with inherited breast and ovarian cancer susceptibility syndrome) and somatic inactivation mechanisms. BRCA-mutant tumors are highly sensitive to genotoxic chemotherapy (Mylavarapu et al., 2018). More recently, targeted therapies for BRCA-mutant tumors have been developed, relying on drugs inhibiting PARP1, which render cells hyper-reliant on the BRCA pathway (Bryant et al., 2005; Farmer et al., 2005; Jackson and Moldovan, 2022). Treatment of BRCA-mutant cells with PARP inhibitors also results in generation of ssDNA gaps, reversed forks prone to degradation, and DSBs (Bryant et al., 2005; Cong et al., 2021a; Farmer et al., 2005; Ray Chaudhuri et al., 2016). The identity of the relevant DNA lesion sensitizing BRCA-deficient cells has been extensively studied, since this has major implications on cancer therapy: providing a biomarker for predicting treatment efficacy, as well as an opportunity for enhancing cancer therapy using inhibitors of mechanisms involved in repairing that specific lesion. Historically, the role of BRCA in recombination-mediated DSB repair has been considered as the critical function of chemotherapy resistance. More recently, its fork protection role was proposed (Bhat and Cortez, 2018; Guillemette et al., 2015; Quinet et al., 2017; Ray Chaudhuri et al., 2016; Taglialatela et al., 2017; Thakar and Moldovan, 2021). Finally, recent work highlighted gap suppression as better correlating with chemotherapy response in particular genetic backgrounds (Berti et al., 2013; Cantor, 2021; Cong et al., 2021b; Dhoonmoon et al., 2022; Jackson et al., 2021; Jackson and Moldovan, 2022; Kang et al., 2021; Panzarino et al., 2021; Quinet et al., 2020; Ray Chaudhuri et al., 2012; Simoneau et al., 2021; Thakar et al., 2022; Thakar et al., 2020; Thakar and Moldovan, 2021; Tirman et al., 2021). However, experiments using separation of function BRCA2 mutants which are proficient for HR but defective in fork protection and gap suppression, suggest that BRCA2 promotes therapy resistance primarily through HR (Feng and Jasin, 2017; Lim et al., 2024).

Potentially reconciling these observations, it was recently shown that ssDNA gaps generate DSBs. Various mechanisms were proposed to explain the etiology of ssDNA gap derived DSBs, including induction of apoptosis (Panzarino et al., 2021) and run-off of the DNA synthesis machinery using the gapped strand as a template in the next cell cycle (Simoneau et al., 2021). To further investigate this, we recently employed PRIMPOL overexpression as a tool to enhance ssDNA gap accumulation in an otherwise DNA repair-proficient genetic background (Nusawardhana et al., 2024). This allowed us to investigate the processing of ssDNA gaps under normal conditions. We found that ssDNA gaps are expanded bidirectionally by the exonuclease activities of MRE11 and EXO1, which have opposing activities. Subsequently, the endonuclease activity of MRE11 cleaves the parental strand at the extended ssDNA gap region, generating a DSB. We also showed that TLS-mediated lesion bypass suppresses this nucleolytic processing, presumably through filling the gaps before the nucleases have the opportunity to engage. Overall, this study indicated that, at least under certain conditions, DSBs are directly generated from ssDNA gaps through nucleolytic processing. However, if this processing is enough to cause chemosensitivity is unclear.

Here, we show that, in DNA repair-proficient backgrounds, ssDNA gap accumulation by itself is not enough to cause chemotherapy sensitivity. Through a series of CRISPR-mediated genome-wide genetic knockout screens in PRIMPOL-overexpressing cells, we identify a pathway centered on the helicase HELQ factors which regulates the cytotoxicity of cisplatin-induced ssDNA gaps. We moreover identify RAD52 as a mediator of this pathway, and show that RAD52 promotes ssDNA gap accumulation through a BRCA-mediated mechanism. Our work identified the HELQ-RAD52-BRCA axis as a regulator of ssDNA gap processing, shedding light on the mechanisms of cellular sensitivity in cancer therapy.

## Results

### PRIMPOL overexpression causes ssDNA gap accumulation without cisplatin sensitization

PRIMPOL overexpression was previously shown to result in formation of ssDNA gaps in response to replication stress (Quinet et al., 2020; Tirman et al., 2021). We recently generated PRIMPOL-overexpressing HeLa and U2OS cells (Figure 1A) (Nusawardhana et al., 2024). In line with these previous reports, PRIMPOL-overexpressing cells showed increased ssDNA gap formation in response to multiple replication stress-inducing agents, including cisplatin and hydroxyurea (HU), as well as the PARP1 inhibitor olaparib (Figure 1B-E, Supplementary Figure S1A-H). Accumulation of ssDNA gaps was detected, in both HeLa and U2OS cells, using two different assays previously employed by us and others to measure nascent strand gaps backgrounds (Berti et al., 2013; Cantor, 2021; Cong et al., 2021b; Dhoonmoon et al., 2022; Jackson et al., 2021; Jackson and Moldovan, 2022; Kang et al., 2021; Panzarino et al., 2021; Quinet et al., 2020; Ray Chaudhuri et al., 2012; Simoneau et al., 2021; Thakar et al., 2022; Thakar et al., 2020; Thakar and Moldovan, 2021; Tirman et al., 2021), namely the BrdU alkaline comet assay (Figure 1B,C; Supplementary Figure S1A-D) and the S1 nuclease DNA fiber assay (Figure 1D,E; Supplementary Figure S1E-H).

**Figure 1.**
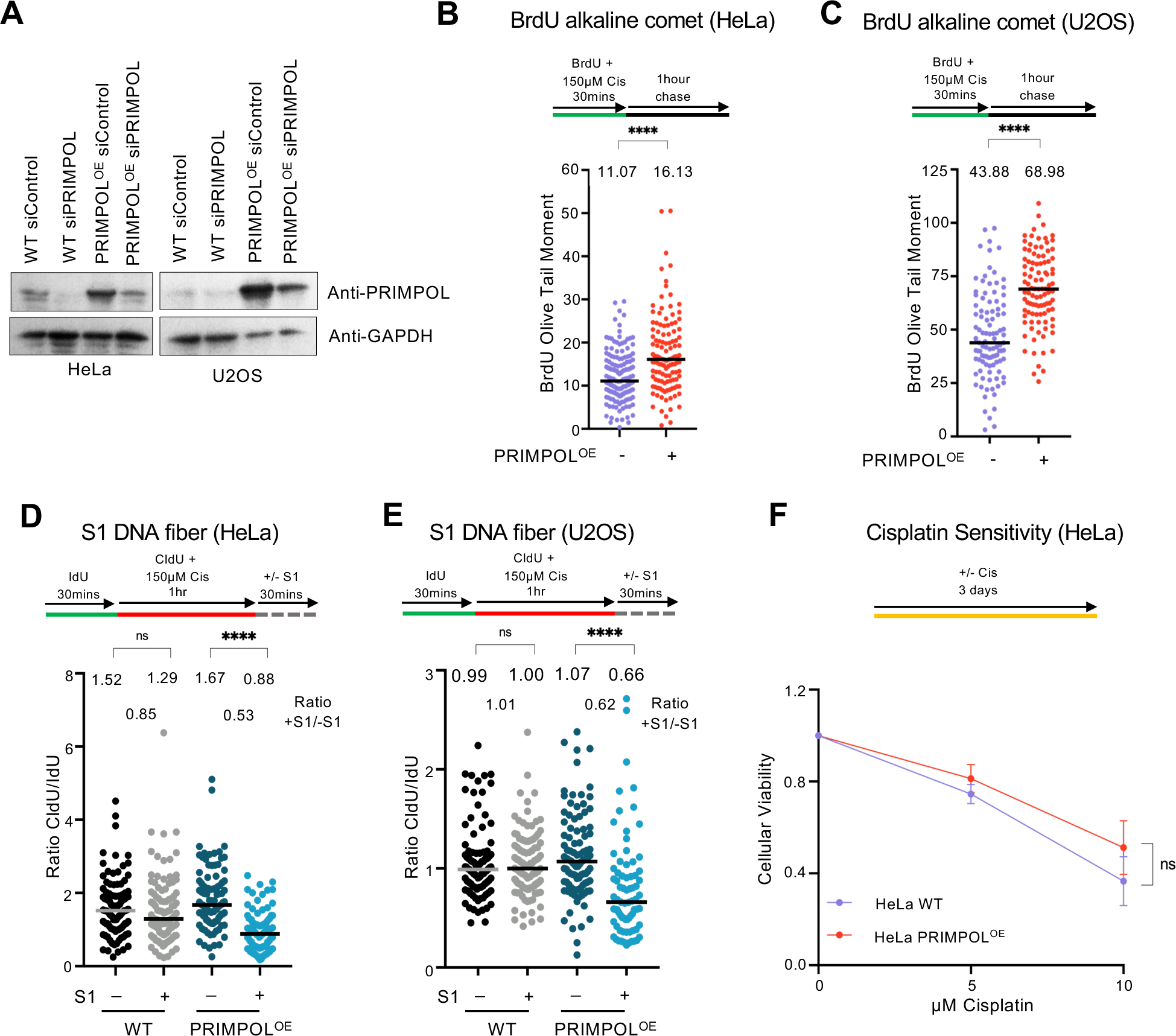
PRIMPOL-overexpressing cells accumulate cisplatin-induced ssDNA gaps but are not sensitive to cisplatin. **A.** Western blots showing PRIMPOL overexpression in HeLa and U2OS cells. **B,C.** BrdU alkaline comet assays showing that PRIMPOL overexpression in HeLa (**B**) and U2OS (**C**) cells causes accumulation of replication-associated ssDNA gaps upon treatment with 150µM cisplatin. At least 95 nuclei were quantified for each condition. The median values are marked on the graph and listed at the top. Asterisks indicate statistical significance (Mann-Whitney, two-tailed). Schematic representations of the assay conditions are shown at the top. **D,E.** S1 nuclease DNA fiber combing assays showing that PRIMPOL overexpression in HeLa (**D**) and U2OS (**E**) cells causes accumulation of nascent strand ssDNA gaps upon treatment with 150µM cisplatin. The ratio of CldU to IdU tract lengths is presented, with the median values marked on the graphs and listed at the top. At least 75 tracts were quantified for each sample. Asterisks indicate statistical significance (Mann-Whitney, two-tailed). Schematic representations of the assay conditions are shown at the top. **F**. Cellular viability assays showing that PRIMPOL overexpression in HeLa cells does not cause cisplatin sensitivity. The average of three independent experiments, with standard deviations indicated as error bars, is shown. Asterisks indicate statistical significance (two-way ANOVA).

Previously, accumulation of ssDNA gaps in certain genetic backgrounds, particularly those with BRCA mutations, were shown to correlate with chemosensitivity (Berti et al., 2013; Cantor, 2021; Cong et al., 2021b; Dhoonmoon et al., 2022; Jackson et al., 2021; Jackson and Moldovan, 2022; Kang et al., 2021; Panzarino et al., 2021; Quinet et al., 2020; Ray Chaudhuri et al., 2012; Simoneau et al., 2021; Thakar et al., 2022; Thakar et al., 2020; Thakar and Moldovan, 2021; Tirman et al., 2021). On the other hand, recent findings suggest that chemosensitivity caused by BRCA2 inactivation mainly reflects the role of BRCA2 in HR, rather than its roles in ssDNA gap suppression and fork protection (Feng and Jasin, 2017; Lim et al., 2024). We observed that, unlike BRCA-deficient cells, PRIMPOL-overexpressing cells are not sensitive to cisplatin (Figure 1F) even though, as shown above, they accumulate cisplatin-induced ssDNA gaps. These findings suggest that gap accumulation may cause chemosensitivity only in certain genetic backgrounds.

### CRISPR screens reveal factors which connect ssDNA gap accumulation to cisplatin sensitivity

To better understand how ssDNA gaps may cause cellular sensitivity, we thought to identify factors which, when inactivated, induce cisplatin sensitivity specifically in PRIMPOL-overexpressing cells, but not in control cells. We hypothesized that these factors are involved in suppressing the processing of ssDNA gaps into cytotoxic structures. To this end, we performed a series of genome-wide CRISPR genetic screens in PRIMPOL-overexpressing and control (empty vector, or EV) HeLa and U2OS cells (Figure 2A). We infected cells with the Brunello genome-wide CRISPR-knockout lentiviral library (Doench et al., 2016), which targets 19,114 human genes with an average of 4 guide RNAs (gRNAs) for each gene, for a total of 76,441 unique gRNAs. After selection, taking care to maintain 250x fold library coverage (equivalent to 20 million cells) at all times, we treated library-infected cells with 0.5µM cisplatin for 14days, splitting cells every 3 days with cisplatin added freshly every time. This cisplatin treatment resulted in ∼30% loss of viability at each splitting time compared to untreated cells. Cells were then collected, and genomic DNA was extracted. The gRNA region was amplified by PCR and identified by Illumina sequencing. Bioinformatic analyses using the MAGeCK algorithm (Li et al., 2014) were used to generate ranking lists of genes which were lost in cisplatin-treated cells compared to untreated cells (Supplementary Tables S1-S4). This represents genes which, when inactivated, result in increased cell death in cisplatin-treated cells compared to untreated cells.

**Figure 2.**
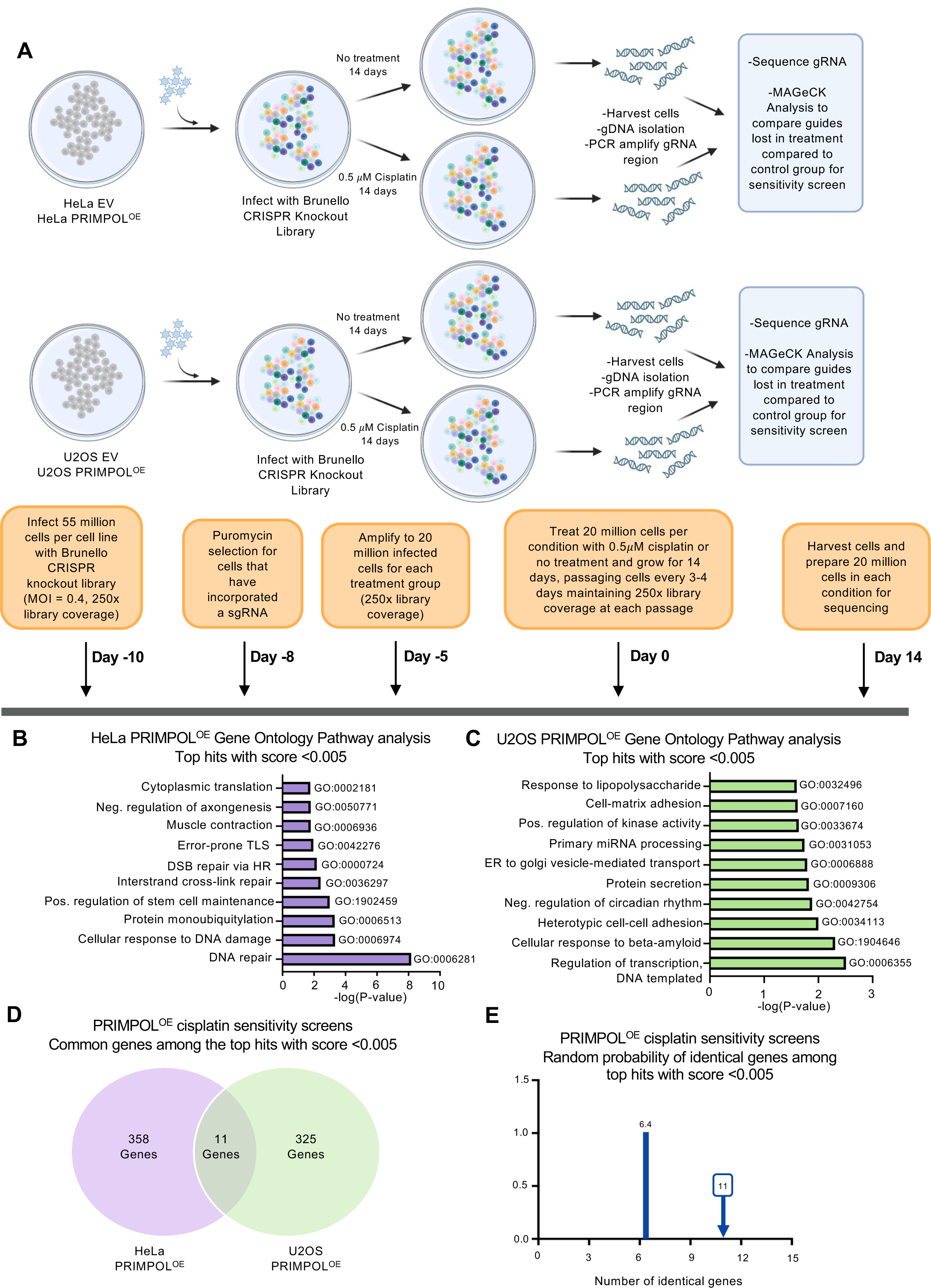
Genome-wide CRISPR knockout screens for cisplatin sensitization of PRIMPOL-overexpressing cells. **A.** Overview of the CRISPR knockout screens to identify genes that are specifically required for cisplatin sensitivity in PRIMPOL-overexpressing cells, but not in control cells. **B,C.** Biological pathway analyses using Gene Ontology analyses of the top hits with MAGeCK score lower than 0.005 which cause cisplatin sensitivity in PRIMPOL-overexpressing HeLa (**B**) and U2OS (**C**) cells. GO_BP terms with negative logP greater than 1.75 (**B**) and 1.60 (**C**) are presented. **D.** Diagram showing the overlap of identical genes within the top hits with MAGeCK score lower than 0.005 which cause cisplatin sensitivity in PRIMPOL-overexpressing HeLa and U2OS cells. **E.** The number of common genes within the top hits with MAGeCK score lower than 0.005 which cause cisplatin sensitivity in PRIMPOL-overexpressing HeLa and U2OS cells (namely 11) is slightly higher than the random probability of identical hits, which is 6.4.

Somewhat surprisingly, we observed differences between the results of the HeLa and U2OS screens, when comparing the genes which are lost in cisplatin-treated PRIMPOL-overexpressing (PRIMPOL^OE^) cells compared to untreated PRIMPOL^OE^ cells. Biological pathway analyses of the top hits of the HeLa-PRIMPOL^OE^ cisplatin sensitivity screen (369 genes with MAGeCK score lower than 0.005), using both Gene Ontology and KEGG networks, revealed DNA repair processes (including interstrand crosslink repair, homologous recombination and translesion synthesis) as major biological mechanisms providing cisplatin resistance (Figure 2B; Supplementary Figure S2A). This is indeed expected considering the mechanism of action of cisplatin. However, we were unable to recapitulate these findings in the U2OS-PRIMPOL^OE^ cisplatin sensitivity screen, which yielded a variety of biological process (including transcription, cell adhesion, and intracellular transport) as top mechanisms upon pathway analyses of the top hits (336 genes with MAGeCK score lower than 0.005) (Figure 2C; Supplementary Figure S2B).

Reflecting these differences, within the top hits (369 genes in the HeLa-PRIMPOL^OE^ cisplatin sensitivity screen and 336 genes in the U2OS-PRIMPOL^OE^ cisplatin sensitivity screen), only 11 were common (Figure 2D). This is only marginally higher than the random probability of common genes within two datasets of these sizes, which is 6.4 (Figure 2E). These 11 genes belong to diverse biological processes, with two of them being DNA repair factors, namely HELQ and GPN1 (Supplementary Figure S2C).

Since our aim was to identify hits which specifically affect cisplatin sensitivity in PRIMPOL-overexpressing compared to control (EV) cells, we performed a similar analysis for the HeLa-EV and U2OS-EV cisplatin sensitivity screens. We found that the results of the HeLa and U2OS screens had reduced overlap in this setup as well. When comparing the genes which are lost in cisplatin-treated EV cells compared to untreated EV cells, out of the top hits with MAGeCK score lower than 0.005 (331 genes in the HeLa-EV screen and 350 genes in the U2OS-EV screen), only 10 were common between the HeLa-EV and U2OS-EV screens (Supplementary Figure S3A,B). This is only marginally higher than the random probability of common genes within two datasets of these sizes, which is 5.9 (Supplementary Figure S3C).

We speculate that the differences between the HeLa and U2OS screen results may be caused by the p53 pathway status, which was previously shown to influence the quality of CRISPR screen results (Bowden et al., 2020; Brown et al., 2019; Haapaniemi et al., 2018; Ihry et al., 2018). Indeed, HeLa cells have a deficient p53 pathway (Matlashewski et al., 1986), which may improve the quality of CRISPR screen results, while U2OS have an active p53 pathway (Allan and Fried, 1999). With this in mind, we focused on identifying hits which were present in both HeLa and U2OS screens.

To this end, we searched in our datasets for genes which were top hits in both the HeLa-PRIMPOL^OE^ and U2OS-PRIMPOL^OE^ cisplatin sensitivity screens, but not in the HeLa-EV or the U2OS-EV cisplatin sensitivity screens. We focused on DNA repair genes which ranked within the top hits (MAGeCK score lower than 0.005) in the PRIMPOL^OE^ cisplatin sensitivity screens (Figure 3A). When listing the ranks of these genes, and comparing the cisplatin treatment to no treatment for each cell line (HeLa-PRIMPOL^OE^, HeLa-EV, U2OS-PRIMPOL^OE^, and U2OS-EV), we noticed that only HELQ fits our criteria: it was a top hit in both HeLa-PRIMPOL^OE^ and U2OS-PRIMPOL^OE^ cisplatin sensitivity screens, but not in the HeLa-EV and U2OS-EV cisplatin sensitivity screens (Figure 3A,B). This suggests that loss of HELQ causes cisplatin sensitivity specifically in PRIMPOL-overexpressing cells compared to cells expressing endogenous levels of PRIMPOL.

**Figure 3.**
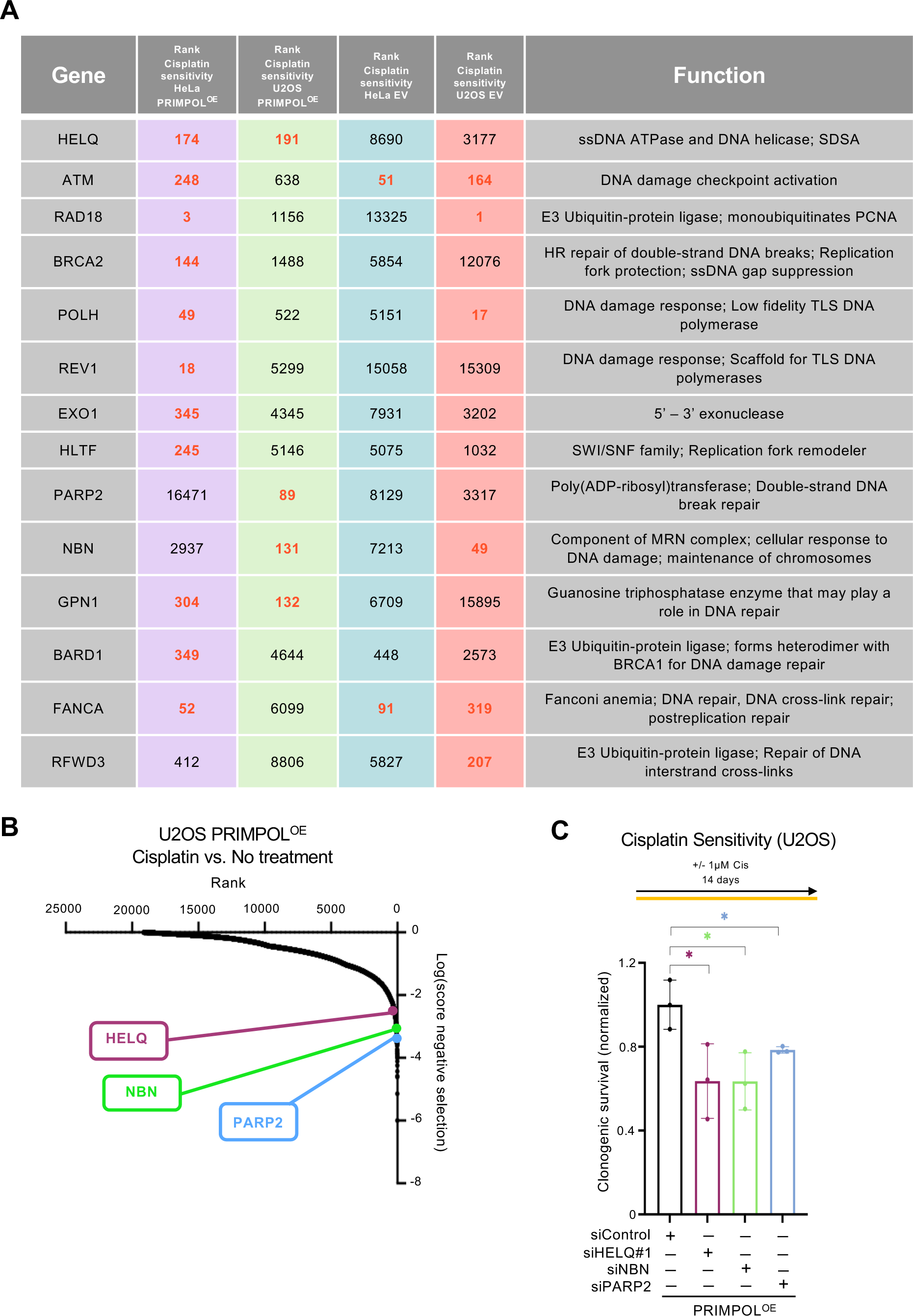
Identification of HELQ as a factor required for suppressing the cisplatin sensitivity of PRIMPOL-overexpressing cells. **A.** Table showing all screens ranking, and the biological functions of the DNA repair top hits in the PRIMPOL^OE^ cisplatin sensitivity screens. The ranks indicated in red represent top ranks in the respective screens (MAGeCK score lower than 0.005). **B.** Scatterplot showing the results of genome-wide CRISPR knockout screens to identify genes that are specifically required for cisplatin sensitivity in PRIMPOL-overexpressing U2OS cells. Three top hits chosen for validation are indicated. **C.** Clonogenic survival assay showing that depletion of HELQ, NBN and PARP2 increases the sensitivity of PRIMPOL-overexpressing U2OS cells to cisplatin treatment (1µM for 14 days). The sensitivity is presented normalized to HeLa-EV control cells. The average of three independent experiments, with standard deviations indicated as error bars, is shown. Asterisks indicate statistical significance (t-test unpaired).

### Loss of HELQ sensitizes PRIMPOL-overexpressing cells to cisplatin and enhances the accumulation of ssDNA gaps and DSBs in these cells

HELQ is a helicase with antagonizing DNA unwinding and strand annealing activities (Anand et al., 2022; Marini and Wood, 2002; Tafel et al., 2011).

In line with the screen results, siRNA-mediated depletion of HELQ resulted in increased cisplatin sensitivity in PRIMPOL-overexpressing cells compared to control cells, as measured using clonogenic assays (Figure 3C). In addition, two other DNA repair genes which were top hits in the U2OS-PRIMPOL^OE^ cisplatin screen, namely PARP2 and NBN, were also validated in U2OS cells (Figure 3B,C). We next thought to investigate if this increased sensitivity is connected to ssDNA gaps. Since HELQ was the only top hit in both HeLa and U2OS screens, we focused our initial investigations on this factor. In both BrdU alkaline comet assays and S1 DNA fiber combing assays, depletion of HELQ did not impact cisplatin-induced ssDNA gap formation in wildtype cells. However, HELQ depletion further exacerbated the increased ssDNA gap accumulation in PRIMPOL-overexpressing cells upon cisplatin treatment. Similar findings were observed in both HeLa and U2OS cells (Figure 4A-D).

**Figure 4.**
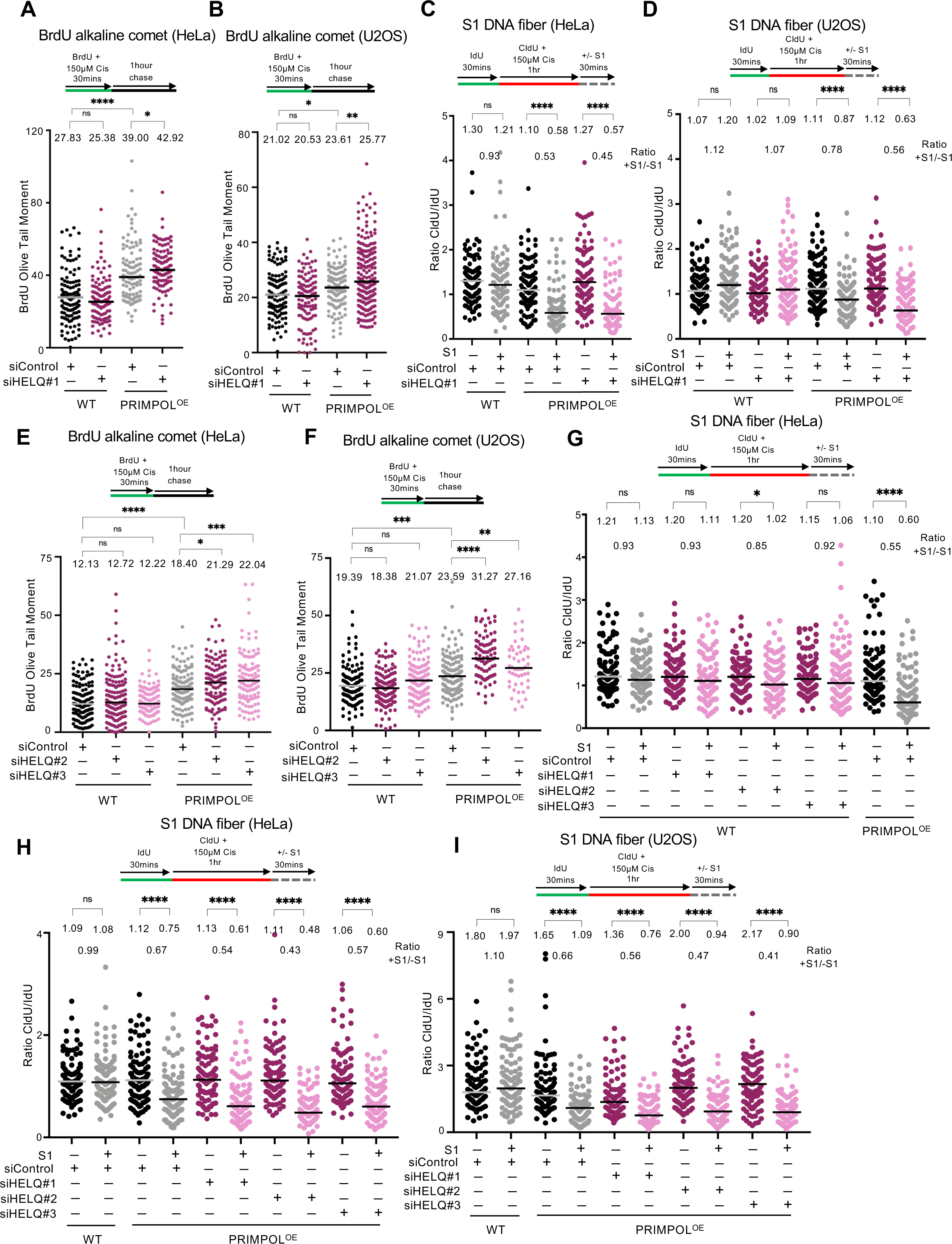
Loss of HELQ increases ssDNA gap accumulation in PRIMPOL-overexpressing cells. **A,B.** BrdU alkaline comet assays showing that HELQ depletion in PRIMPOL-overexpressing HeLa (**A**) and U2OS (**B**) cells causes accumulation of replication-associated ssDNA gaps upon treatment with 150µM cisplatin. At least 100 nuclei were quantified for each condition. The median values are marked on the graph and listed at the top. Asterisks indicate statistical significance (Mann-Whitney, two-tailed). Schematic representations of the assay conditions are shown at the top. **C,D.** S1 nuclease DNA fiber combing assays showing that HELQ depletion in PRIMPOL-overexpressing HeLa (**C**) and U2OS (**D**) cells causes accumulation of nascent strand ssDNA gaps upon treatment with 150µM cisplatin, as indicated by a decrease in the +S1/−S1 ratios in the HELQ-depleted compared to control-depleted PRIMPOL-overexpressing cells. The ratio of CldU to IdU tract lengths is presented, with the median values marked on the graphs and listed at the top. The +S1/−S1 ratios of the median values are also presented. At least 100 tracts were quantified for each sample. Asterisks indicate statistical significance (Mann-Whitney, two-tailed). Schematic representations of the assay conditions are shown at the top. **E,F.** BrdU alkaline comet assays showing that HELQ depletion using additional siRNA oligonucleotides in PRIMPOL-overexpressing HeLa (**E**) and U2OS (**F**) cells causes accumulation of replication-associated ssDNA gaps upon treatment with 150µM cisplatin. At least 50 nuclei were quantified for each condition. The median values are marked on the graph and listed at the top. Asterisks indicate statistical significance (Mann-Whitney, two-tailed). Schematic representations of the assay conditions are shown at the top. **G-I.** S1 nuclease DNA fiber combing assays showing that HELQ depletion does not cause ssDNA gap accumulation in control HeLa cells (**G**), but increases ssDNA gap accumulation in PRIMPOL-overexpressing HeLa (**H**) and U2OS (**I**) cells upon treatment with 150µM cisplatin, as indicated by a decrease in the +S1/−S1 ratios in the HELQ-depleted compared to control-depleted PRIMPOL-overexpressing cells. The ratio of CldU to IdU tract lengths is presented, with the median values marked on the graphs and listed at the top. The +S1/−S1 ratios of the median values are also presented. At least 100 tracts were quantified for each sample. Asterisks indicate statistical significance (Mann-Whitney, two-tailed). Schematic representations of the assay conditions are shown at the top. Western blots confirming HELQ depletion are shown in Supplementary Figure S4A.

In order to rule out off-target effects of the siRNA oligonucleotide employed, we repeated these analyses using 2 additional siRNA oligonucleotides targeting HELQ. Depletion of HELQ using these oligonucleotides confirmed the initial results. Loss of HELQ increased cisplatin-induced ssDNA gaps in PRIMPOL-overexpressing HeLa or U2OS cells but not in control cells, as measured using the BrdU alkaline comet assay (Figure 4E,F; Supplementary Figure S4A). HELQ depletion by multiple siRNA oligonucleotides also did not increase gap formation in wildtype control cells as measured by the S1 nuclease DNA fiber combing assay (Figure 4G). Since PRIMPOL-overexpressing cells already showed significant ssDNA gap accumulation, a further increase in this accumulation can be difficult to observe in the S1 nuclease fiber combing assay. We thus calculated the +S1/−S1 ratios (median of CldU/IdU ratios of S1-treated samples divided to the median of CldU/IdU ratios of non-S1-treated samples) for each condition. A decrease in the +S1/−S1 ratios in the HELQ-depleted PRIMPOL^OE^ cells compared to mock depleted (siControl) PRIMPOL^OE^ cells was observed in both HeLa and U2OS cells (Figure 4H,I). This indicates that HELQ depletion further increases gap accumulation in PRIMPOL-overexpressing cells, thus confirming the BrdU alkaline comet assay results.

To further confirm these results in a separate experimental system, we knocked out HELQ in HeLa cells using CRISPR/Cas9. Two independent HELQ-knockout clones were obtained. We then introduced the PRIMPOL-overexpressing construct (or empty vector as control) in these cells (Figure 5A). Clonogenic cisplatin sensitivity experiments indicated that both HELQ-knockout PRIMPOL-overexpression lines have increased cisplatin sensitivity compared to the wildtype (HELQ-proficient) PRIMPOL-overexpressing cells (Figure 5B). These findings further validate the CRISPR screen results presented in Figure 1. We then measured cisplatin-induced ssDNA gap accumulation in these cells. Similar to the results using HELQ depletion by siRNA, the +S1/−S1 ratios were lower in both HELQ-knockout PRIMPOL-overexpressing cell lines compared to HELQ-proficient PRIMPOL-overexpression cells (Figure 5C).

**Figure 5.**
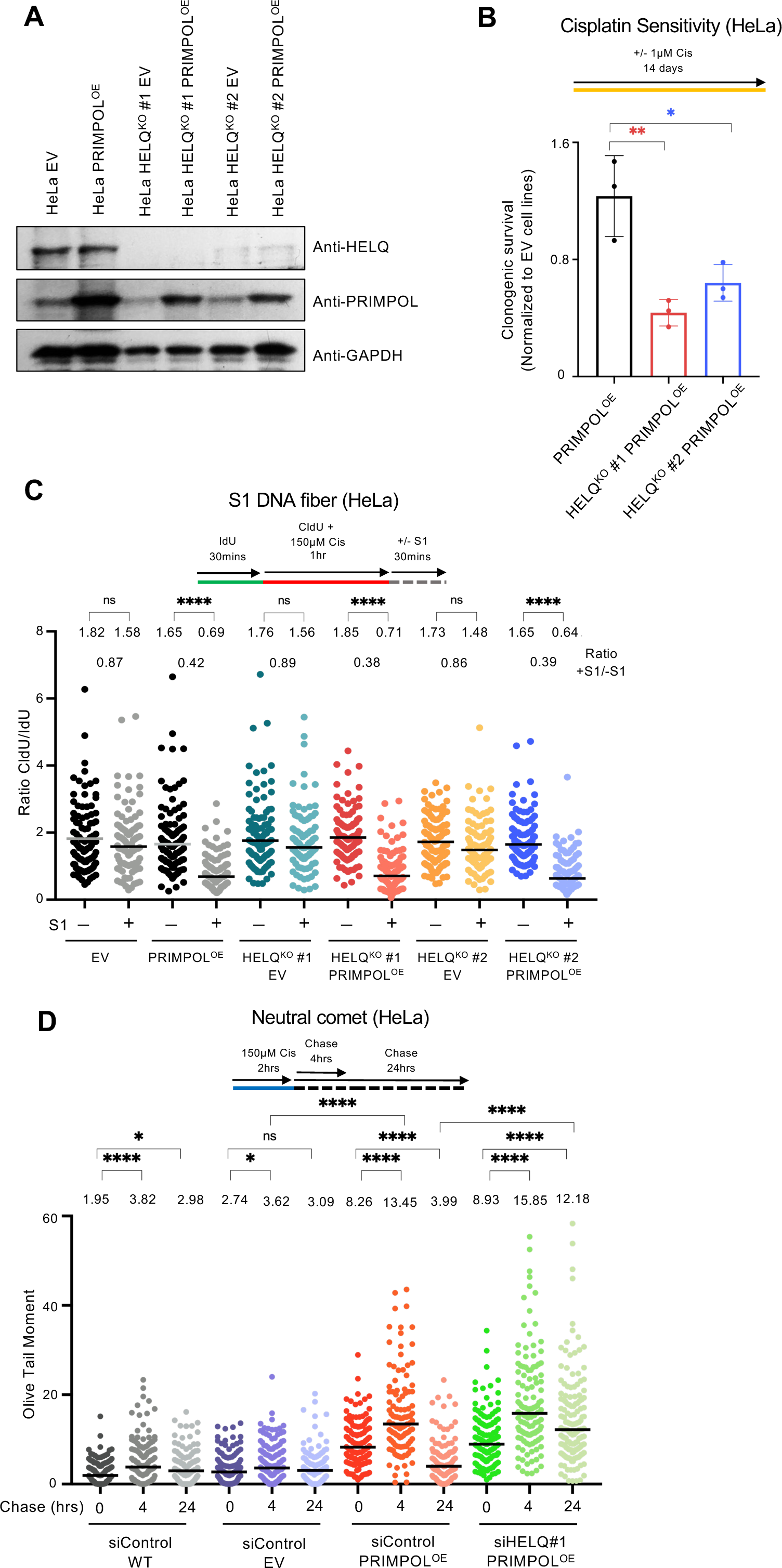
HELQ controls genomic stability of PRIMPOL-overexpressing cells. **A.** Western blots showing HELQ knockout in HeLa cells and PRIMPOL overexpression in these cells. **B.** Clonogenic survival assay showing that HELQ knockout specifically increases the sensitivity of PRIMPOL-overexpressing HeLa cells to cisplatin treatment (1µM for 14 days). The sensitivity is presented normalized to HeLa-EV control cells. The average of three independent experiments, with standard deviations indicated as error bars, is shown. Asterisks indicate statistical significance (t-test, unpaired). **C.** S1 nuclease DNA fiber combing assay showing that HELQ knockout does not cause ssDNA gap accumulation in control HeLa-EV (empty vector) cells, but increases ssDNA gap accumulation in PRIMPOL-overexpressing HeLa cells upon treatment with 150µM cisplatin, as indicated by a decrease in the +S1/−S1 ratios. The ratio of CldU to IdU tract lengths is presented, with the median values marked on the graph and listed at the top. The +S1/−S1 ratios of the median values are also presented. At least 100 tracts were quantified for each sample. Asterisks indicate statistical significance (Mann-Whitney, two-tailed). A schematic representation of the assay conditions is shown at the top. **D**. Neutral comet assays showing that treatment with 150µM cisplatin for 2 hours causes accumulation of DSBs in PRIMPOL-overexpressing HeLa cells 4 hours later, which are repaired after 24 hours in HELQ-proficient, but not in HELQ siRNA-depleted cells. At least 110 comets were quantified for each sample. The median values are marked on the graph, and asterisks indicate statistical significance (Mann-Whitney, two-tailed).

The increase in ssDNA gap formation upon HELQ deficiency in PRIMPOL-overexpression cells seems to be minimal, at least in the S1 nuclease DNA fiber combing assay. While this may reflect a limitation of the assay, it does raise the question of whether such a minimal increase in ssDNA gap accumulation can account for the significant increase in cisplatin sensitivity observed under these conditions. To potentially explain this, we reasoned that the increase in ssDNA gaps upon HELQ depletion may generate a more robust increase in DSBs. Indeed, we recently showed that cisplatin-induced ssDNA gaps accumulating in PRIMPOL-overexpressing cells are ultimately converted into double stranded DNA breaks (DSBs); this conversion takes place within 2 hrs of cisplatin exposure (Nusawardhana et al., 2024). However, since PRIMPOL-overexpressing cells are not cisplatin-sensitive even though they accumulate ssDNA gaps and subsequently DSBs, we reasoned that these DSBs are eventually repaired, since the cells are BRCA-proficient. To test this, we employed the neutral comet assay to monitor cisplatin-induced DSB formation following cisplatin removal (Figure 5D). As we previously showed, after two hours of cisplatin treatment the PRIMPOL-overexpressing cells have more DSBs than control (EV) cells. We then removed cisplatin-containing media and grew cells in fresh cisplatin-free media. There were no changes in DSB formation in control cells. In contrast, in PRIMPOL-overexpressing cells we detected an increase in DSB formation after 4hrs of growth in fresh media, indicating that DSBs continue to form, presumably from processing of ssDNA gaps which were formed during the cisplatin treatment. However, 24h later the amount of DSBs was reduced, indicating that these gaps were repaired, and potentially explaining why PRIMPOL-overexpressing cells are not cisplatin sensitive even though they accumulate cisplatin-induced ssDNA gaps. We then examined the impact of HELQ depletion on the dynamics of ssDNA gaps-derived DSBs. In contrast to HELQ-proficient (siControl) PRIMPOL-overexpressing cells, HELQ-depleted PRIMPOL-overexpressing cells showed persistent DSB accumulation at 24h following removal of cisplatin (Figure 5D). These findings suggest that, in PRIMPOL-overexpressing cells, HELQ is necessary for the repair of DSBs derived from cisplatin-induced ssDNA gaps.

### RAD52 promotes ssDNA gap accumulation in PRIMPOL-overexpressing cells but not in BRCA-deficient cells

Recent studies have shown that HELQ interacts with the POLD3 subunit of the DNA replicative polymerase Polδ to suppress DNA synthesis during break-induced replication (BIR) and promote DNA strand annealing (He et al., 2023). We thought to investigate if loss of POLD3 has a similar impact as HELQ on ssDNA gap accumulation in PRIMPOL-overexpressing cells. We also extended our analyses to RAD52, another strand annealing factor that participates in BIR upstream of POLD3 (Rossi et al., 2021). We thus depleted POLD3 or RAD52 in PRIMPOL-overexpressing cells and measured ssDNA gap accumulation upon cisplatin treatment. We first performed BrdU alkaline comet assays. Depletion of POLD3, with two different siRNA oligonucleotides, did not affect ssDNA gap accumulation in either PRIMPOL-overexpressing cells or control cells (Figure 6A; Supplementary Figure S4B). These results argue against POLD3 and its interaction with HELQ being involved in the HELQ-mediated gap expansion described above in PRIMPOL-overexpressing cells.

**Figure 6.**
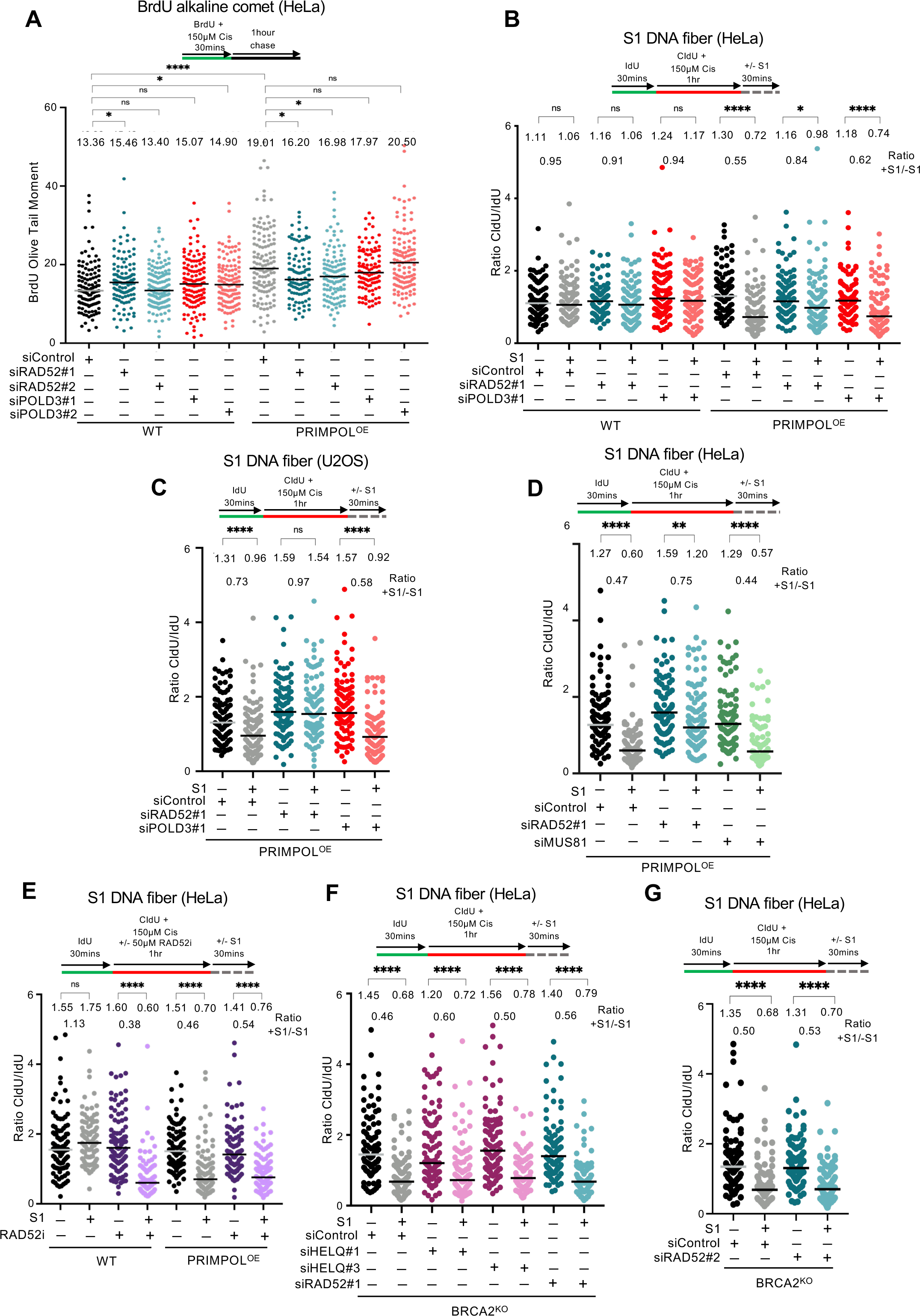
Loss of RAD52 suppresses ssDNA gap accumulation in PRIMPOL-overexpressing cells but not in BRCA-deficient cells. **A.** BrdU alkaline comet assays showing that RAD52 depletion reduces ssDNA gap accumulation upon treatment with 150µM cisplatin in PRIMPOL-overexpressing HeLa cells but not in control HeLa cells, while POLD3 depletion does not affect it. At least 100 nuclei were quantified for each condition. The median values are marked on the graph and listed at the top. Asterisks indicate statistical significance (Mann-Whitney, two-tailed). A schematic representation of the assay conditions is shown at the top. Western blots confirming depletion of RAD52 and of POLD3 are shown in Supplementary Figure S4B,C. **B,C.** S1 nuclease DNA fiber combing assays showing that RAD52 depletion suppresses ssDNA gap accumulation upon treatment with 150µM cisplatin in PRIMPOL-overexpressing HeLa (**B**) and U2OS (**C**) cells but not in control cells, while POLD3 depletion does not affect it. The ratio of CldU to IdU tract lengths is presented, with the median values marked on the graphs and listed at the top. The +S1/−S1 ratios of the median values are also presented. At least 65 tracts were quantified for each sample. Asterisks indicate statistical significance (Mann-Whitney, two-tailed). Schematic representations of the assay conditions are shown at the top. **D.** S1 nuclease DNA fiber combing assays showing that RAD52 depletion suppresses ssDNA gap accumulation upon treatment with 150µM cisplatin in PRIMPOL-overexpressing HeLa cells, while MUS81 depletion does not affect it. The ratio of CldU to IdU tract lengths is presented, with the median values marked on the graphs and listed at the top. The +S1/−S1 ratios of the median values are also presented. At least 70 tracts were quantified for each sample. Asterisks indicate statistical significance (Mann-Whitney, two-tailed). Schematic representations of the assay conditions are shown at the top. Western blots confirming depletion of MUS81 are shown in Supplementary Figure S4D. **E.** S1 nuclease DNA fiber combing assays showing that treatment with the RAD52 inhibitor (-Epigallocatechin) (50µM) causes ssDNA gap accumulation upon exposure to 150µM cisplatin in HeLa cells. The ratio of CldU to IdU tract lengths is presented, with the median values marked on the graphs and listed at the top. The +S1/−S1 ratios of the median values are also presented. At least 95 tracts were quantified for each sample. Asterisks indicate statistical significance (Mann-Whitney, two-tailed). A schematic representation of the assay conditions is shown at the top. **F,G.** S1 nuclease DNA fiber combing assays showing that RAD52 depletion does not impact ssDNA gap accumulation upon treatment with 150µM cisplatin in BRCA2-knockout HeLa cells. The ratio of CldU to IdU tract lengths is presented, with the median values marked on the graphs and listed at the top. The +S1/−S1 ratios of the median values are also presented. At least 100 tracts were quantified for each sample. Asterisks indicate statistical significance (Mann-Whitney, two-tailed). Schematic representations of the assay conditions are shown at the top.

In contrast, we surprisingly observed that RAD52 depletion, with two different siRNA oligonucleotides, suppressed ssDNA gap accumulation in PRIMPOL-overexpressing cells upon cisplatin treatment, while not generally affecting this in control cells (Figure 6A; Supplementary Figure S4C). Similar findings were observed upon ssDNA gap induction by olaparib in PRIMPOL-overexpressing cells, where HELQ depletion increased gap accumulation, POLD3 depletion did not affect it, and RAD52 depletion decreased it (Supplementary Figure S5A).

We next sought to validate these studies using the S1 nuclease DNA fiber combing assay. Similar to the BrdU alkaline comet assay results, depletion of POLD3 or of RAD52 did not cause ssDNA gap accumulation in cisplatin-treated wildtype cells. In contrast, RAD52 depletion suppressed ssDNA gap accumulation in PRIMPOL-overexpressing cells, while POLD3 did not affect it. Similar results were obtained in both HeLa and U2OS cells (Figure 6B,C; Supplementary Figure S5B,C). Moreover, depletion of MUS81, which participates in BIR downstream of RAD52 and upstream of POLD3 (Rossi et al., 2021), also did not suppress cisplatin-induced ssDNA gap accumulation in PRIMPOL-overexpressing HeLa cells (Figure 6D, Supplementary Figure S4D). Overall, these findings indicate that RAD52 activity promotes ssDNA gap accumulation in PRIMPOL-overexpressing cells, and this does not occur through the role of RAD52 in POLD3/MUS81-mediated BIR.

Recently, RAD52 was proposed to suppress ssDNA gaps generated by Polα in HU-treated wildtype U2OS cells (Di Biagi et al., 2023). In our hands, RAD52 depletion did not have a major effect on ssDNA gap accumulation in cisplatin-treated wildtype cells. In contrast to our experiments using RAD52 depletion, this study employed the RAD52 inhibitor EGC to inactivate its activity. We reasoned that differences between RAD52 depletion and inhibition may account for these discrepancies. Indeed, unlike RAD52 depletion, its inhibition using EGC caused cisplatin-induced ssDNA gap accumulation in wildtype cells (Figure 6E), in line with this previous report. These findings indicate that inhibition of RAD52 by EGC results in additional effects on gap formation compared to the loss of RAD52 protein. We thus decided to employ RAD52 siRNA-mediated depletion rather than RAD52 inhibition for our subsequent studies.

Previous studies showed that BRCA-deficient cells accumulate gaps upon treatment with cisplatin and other replication stress-inducing agents, and this may correlate with their chemosensitivity (Cong et al., 2021a; Hale et al., 2023; Panzarino et al., 2021; Quinet et al., 2021; Quinet et al., 2020; Tirman et al., 2021). The surprising gap suppression effect observed upon RAD52 depletion in PRIMPOL-overexpressing cells prompted us to investigate if a similar effect occurs in BRCA-deficient cells. To this end, we employed the HeLa BRCA2-knockout cells previously generated in our laboratory (Clements et al., 2018) which we recently described to accumulate cisplatin-induced ssDNA gaps (Hale et al., 2023). In contrast to the almost complete suppression of gap formation observed in PRIMPOL-overexpressing cells, RAD52 depletion, using 2 different siRNA oligonucleotides, only had a minimal effect on gap accumulation in BRCA2-knockout cells (Figure 6F,G). Moreover, HELQ depletion also did not affect gap formation in BRCA2-knockout cells (Figure 6F). Overall, these findings indicate that RAD52 promotes the formation of ssDNA gaps generated in PRIMPOL-overexpressing cells, but does not impact gap generation in BRCA-deficient cells.

### RAD52 controls ssDNA gap formation in a HELQ and BRCA-dependent manner

The results presented above indicated that HELQ and RAD52 have opposite effects on ssDNA gap accumulation in PRIMPOL-overexpressing cells. To gain insights into the mechanism behind their roles in ssDNA gap metabolism, we tested for potential genetic interactions between these factors. For this, we made use of the HELQ-knockout PRIMPOL-overexpressing cells described above. In contrast to the suppression described above in HELQ-proficient PRIMPOL-overexpressing cells, RAD52 depletion in HELQ-knockout PRIMPOL-overexpressing cells failed to suppress gap accumulation. Similar findings were observed using both the BrdU alkaline comet and the S1 nuclease DNA fiber combing assays (Figure 7A,B). These results indicate that the presence of HELQ is necessary for the gap suppression observed upon RAD52 depletion in PRIMPOL-overexpressing cells, and suggest that RAD52 promotes ssDNA gap formation in these cells in a manner which requires HELQ.

**Figure 7.**
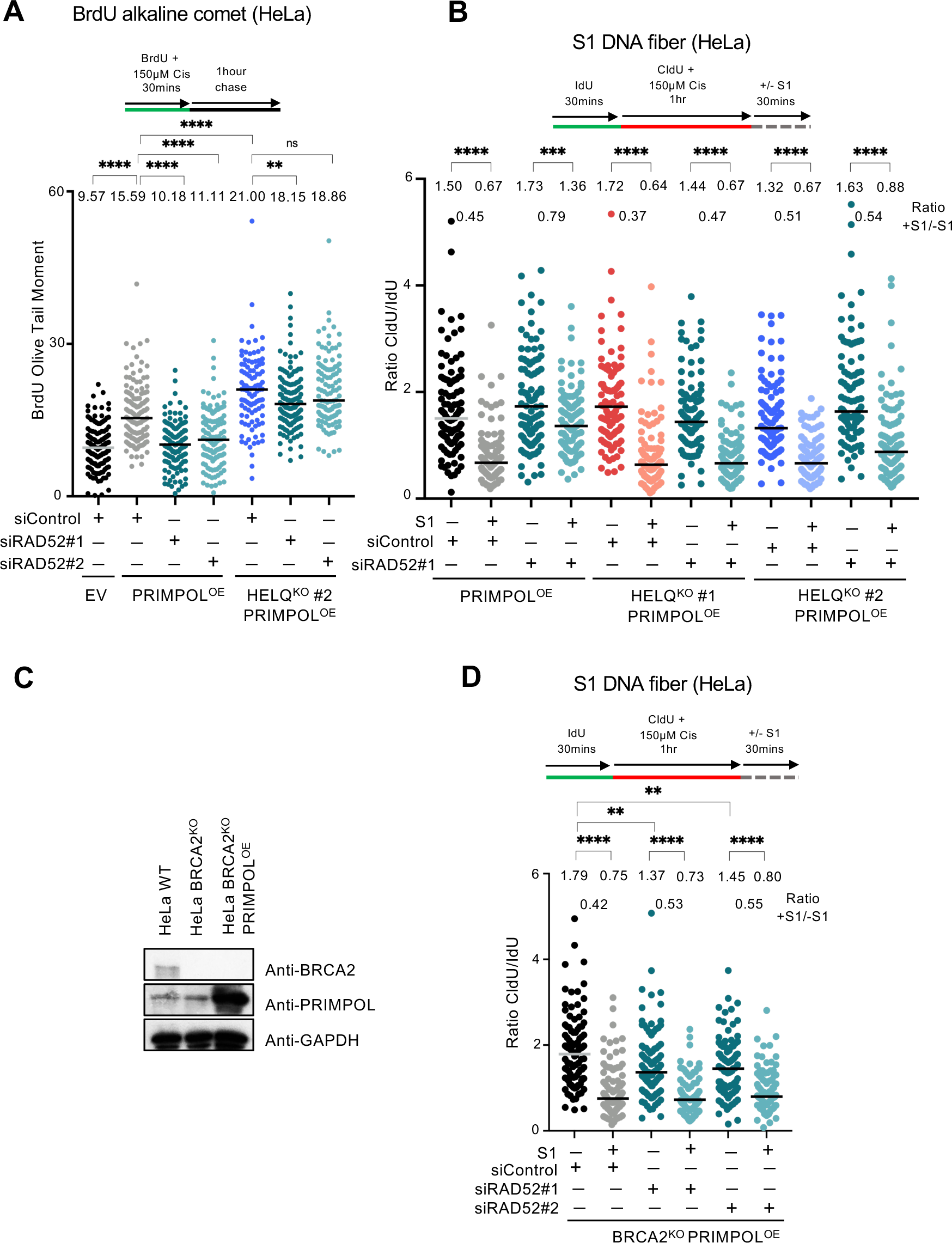
The role of RAD52 in gap formation requires HELQ and the BRCA pathway. **A.** BrdU alkaline comet assays showing that RAD52 depletion suppresses ssDNA gap accumulation in HELQ-proficient, but not in HELQ-knockout PRIMPOL-overexpressing HeLa cells, upon treatment with 150µM cisplatin. At least 100 nuclei were quantified for each condition. The median values are marked on the graph and listed at the top. Asterisks indicate statistical significance (Mann-Whitney, two-tailed). A schematic representation of the assay conditions is shown at the top. **B.** S1 nuclease DNA fiber combing assays showing that RAD52 depletion suppresses ssDNA gap accumulation in HELQ-proficient, but not in HELQ-knockout PRIMPOL-overexpressing HeLa cells, upon treatment with 150µM cisplatin. The ratio of CldU to IdU tract lengths is presented, with the median values marked on the graphs and listed at the top. The +S1/−S1 ratios of the median values are also presented. At least 100 tracts were quantified for each sample. Asterisks indicate statistical significance (Mann-Whitney, two-tailed). A schematic representation of the assay conditions is shown at the top. **C.** Western blots showing PRIMPOL overexpression in BRCA2-knockout HeLa cells. **D.** S1 nuclease DNA fiber combing assays showing that RAD52 depletion does not impact ssDNA gap accumulation upon treatment with 150µM cisplatin in PRIMPOL-overexpressing BRCA2-knockout HeLa cells. The ratio of CldU to IdU tract lengths is presented, with the median values marked on the graphs and listed at the top. The +S1/−S1 ratios of the median values are also presented. At least 100 tracts were quantified for each sample. Asterisks indicate statistical significance (Mann-Whitney, two-tailed). A schematic representation of the assay conditions is shown at the top.

We next investigated if any of the previously-described activities of RAD52 are involved in the function of RAD52 in ssDNA gap metabolism described here. Arrested forks can be restarted by either fork reversal or PRIMPOL repriming. RAD52 was previously shown to suppress fork reversal by antagonizing the recruitment of SMARCAL1 translocase to arrested forks (Honda et al., 2023; Malacaria et al., 2019). An increase in fork reversal upon RAD52 depletion may potentially explain the suppression of ssDNA gap accumulation in PRIMPOL-overexpressing cells described above. However, in our experiments we did not observe a reduction in the CldU/IdU ratios in the non-S1 nuclease-treated samples in RAD52-depleted cells compared to control cells (Figure 6), suggesting that increased fork reversal does not occur under these conditions and thus the suppression of gap accumulation cannot be explained by this.

RAD52 was also previously suggested to participate, directly or indirectly, in the recruitment of the MRE11 nuclease to stalled forks in BRCA-deficient cells (Mijic et al., 2017; Ronson et al., 2023). Recent studies indicated that MRE11 participates in the exonucleolytic expansion of ssDNA gaps in both BRCA-deficient cells as well as in PRIMPOL-overexpressing cells (Hale et al., 2023; Nusawardhana et al., 2024; Tirman et al., 2021). A reduction in MRE11 recruitment to ssDNA gaps may potentially explain the suppression of ssDNA gap accumulation in PRIMPOL-overexpressing cells. However, our findings indicate that loss of RAD52 does not suppress ssDNA gap accumulation in BRCA2-deficient cells (Figure 6), despite the previously-published suppression of MRE11 recruitment in these cells, described above. This argues against a role for MRE11 recruitment in RAD52-dependent ssDNA gap formation observed in PRIMPOL-overexpressing cells.

The fact that RAD52 depletion was not able to suppress gap formation in BRCA2-knockout cells prompted us to investigate if, instead of fork reversal or MRE11 recruitment, the impact of RAD52 on gap accumulation in PRIMPOL-overexpressing cells involves the BRCA pathway. To test this, we overexpressed PRIMPOL in BRCA2-knockout cells (Figure 7C). Unlike the suppression observed in BRCA-proficient PRIMPOL-overexpression cells, depletion of RAD52 was not able to suppress cisplatin-induced ssDNA gap accumulation in BRCA2-knockout PRIMPOL-overexpressing cells (Figure 7D). Similarly, HELQ depletion did not impact gap formation in these cells (Supplementary Figure S5D). Overall, these findings suggest that RAD52 promotes gap formation in PRIMPOL-overexpressing cells through the BRCA pathway.

## Discussion

Our work indicates that RAD52 promotes ssDNA gap accumulation in PRIMPOL-overexpressing cells, but not in BRCA2-deficient cells. Moreover, we show that this role of RAD52 in promoting gap accumulation in PRIMPOL-overexpressing cells does not appear to reflect its previously-described roles in single strand annealing, suppression of fork reversal or recruitment of the MRE11 nuclease. Instead, we show that an intact BRCA pathway is required for this, suggesting that the presence of RAD52 blocks BRCA-mediated gap filling in PRIMPOL-overexpressing cells. We moreover show that this also requires HELQ, perhaps suggesting a role for HELQ in promoting BRCA-mediated gap filling in RAD52-depleted PRIMPOL-overexpressing cells. While it is still unclear how exactly the HELQ-RAD52-BRCA axis regulates gap accumulation in PRIMPOL-overexpressing cells, our results suggest that it involves BRCA-mediated gap filling. While previous work clearly showed that BRCA-deficient cells accumulate ssDNA gaps, how the BRCA pathway suppresses gap formation is still unclear. We speculate that at least one of the mechanisms through which this suppression occurs involves a recombination event with the nascent strand of the sister chromatid (Figure 8A). However, for this event to occur, a displacement (D-) loop needs to be formed on the sister chromatid, upon its invasion by a 3’ end derived from a homologous strand. Most likely, this is the 3’ end of the interrupted nascent strand at the ssDNA gap region, since no other 3’ end exists in this structure. However, this strand is normally annealed to the intact parental strand. We speculate that HELQ, through its helicase activity, may be responsible for unwinding this 3’ end, allowing it to be coated with RAD51 nucleofilaments in a BRCA-catalyzed event. This structure can then invade the sister chromatid and form homology with the intact nascent strand on that chromatid. DNA synthesis extension of the 3’ end ensures replication through the gapped region, and upon re-annealing to the parental strand of the original chromatid, the gap is filled. HELQ activity may also be required at this stage (in addition to or instead of the 3’ end unwinding), since previous studies have shown that HELQ participates in 3’ end extension downstream of D-loop formation, potentially through removing RAD51 from the dsDNA formed at the D-loop, and thus allowing for DNA polymerase engagement (Adelman et al., 2013; Thomas et al., 2022; Ward et al., 2010).

**Figure 8.**
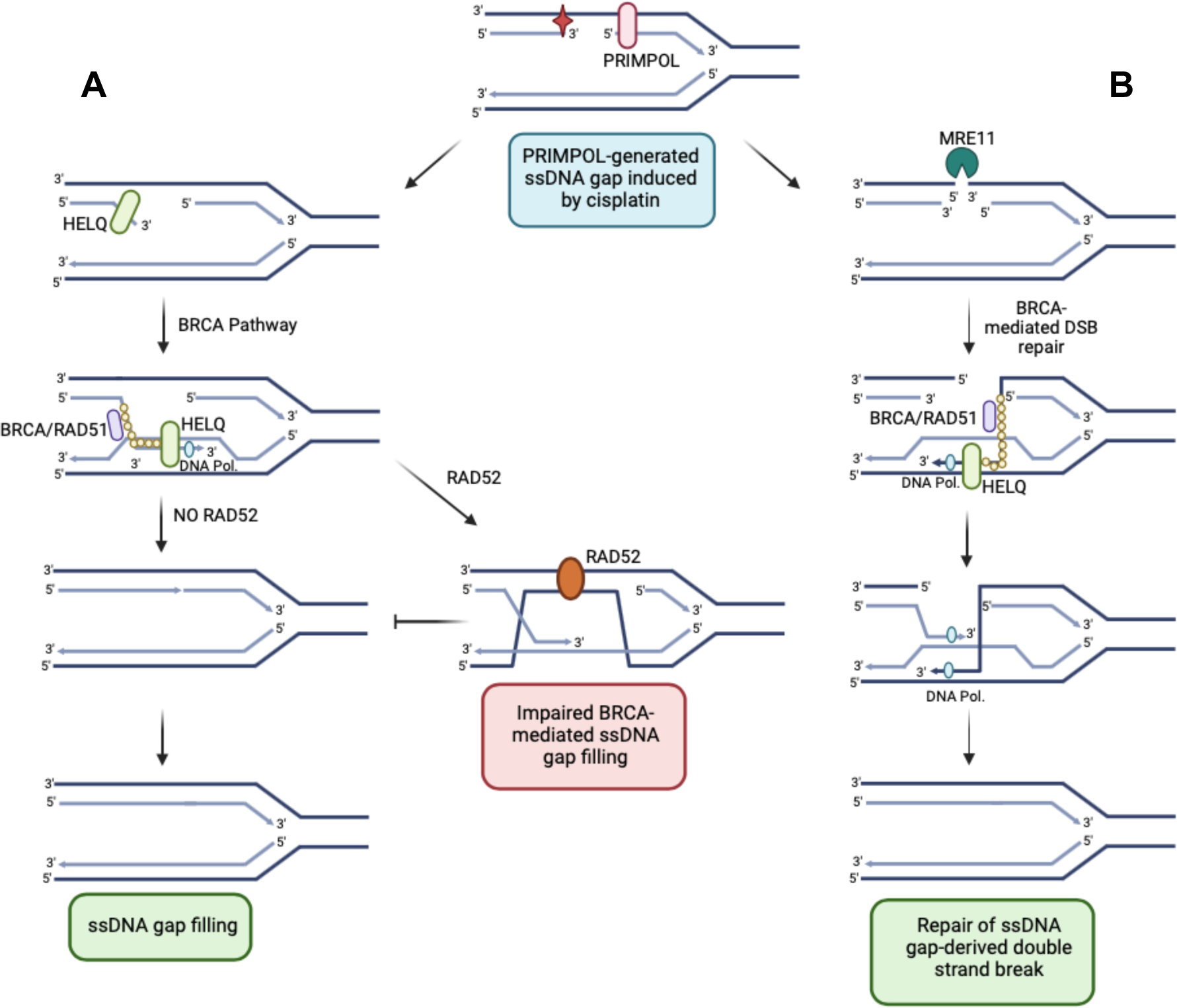
Speculative models for the roles of HELQ and RAD52 in ssDNA gap formation. **A.** PRIMPOL-derived gaps are repaired through BRCA-mediated gap filling. HELQ promotes this process, potentially at multiple steps. In contrast, by catalyzing the annealing of the displaced strand to the intact parental strand on the gapped chromatid, RAD52 interferes with this process. **B**. PRIMPOL-derived gaps can also be repaired by BRCA-mediated homologous recombination upon DSB formation. HELQ promotes this recombination process as well. Created with Biorender.com.

Our work also suggests that RAD52 antagonizes gap filling in this setup. In the HR-mediated repair of DSBs, RAD52 mediates second end capture, in which the third strand in the D-loop structure anneals to the resected DNA on the other end of the DSB, forming a double Holiday junction which allows DNA synthesis to fill the other broken strand (Di Biagi et al., 2023). However, in gap filling, the other strand (in this case the parental strand) is not broken. We speculate that, in this scenario, second strand capture catalyzed by RAD52 results in the formation of a structure which impedes the repair process, since no 3’ end is present in this structure to be extend through DNA synthesis. Thus, RAD52 activity would be toxic to the ssDNA gap filling process in this situation, potentially explaining why loss of RAD52 promotes gap filling in PRIMPOL-overexpressing cells in a manner dependent on HELQ and the BRCA pathway. On the other hand, it was previously proposed that RAD52 competes with BRCA2 binding to resected 3’ ends (Yasuda et al., 2018). It is thus possible that the presence of RAD52 at gap structures may inhibit the recruitment of BRCA2 and subsequent BRCA-mediated gap filling through a mechanism independent of RAD51 loading.

Overall, our work shows that ssDNA gap accumulation is not always associated with chemosensitivity. PRIMPOL-overexpressing cells show cisplatin-induced ssDNA gap accumulation, but not cisplatin sensitivity. HELQ depletion in PRIMPOL-overexpressing cells causes mildly increased ssDNA gap accumulation and sensitizes these cells to cisplatin. While the model presented above may explain the differential impact of HELQ and RAD52 loss on ssDNA gap accumulation, it does not explain why HELQ depletion also potentiates cisplatin-induced DSB accumulation and causes cisplatin sensitivity in PRIMPOL-overexpressing cells. To explain this, we speculate that HELQ also participates in a separate pathway of gap repair. We previously showed that PRIMPOL-overexpressing cells accumulate ssDNA-gap derived DSBs (Nusawardhana et al., 2024). We show here that these DSB are eventually repaired in HELQ-proficient cells, but not in HELQ-deficient cells, suggesting that the loss of HELQ suppresses the repair of ssDNA gap-derived DSBs. In our previous work (Nusawardhana et al., 2024), we showed that the DSBs are formed through the endonucleolytic activity of MRE11 on the parental strand at the ssDNA gap region. Since these DSBs are eventually repaired, we hypothesize that the MRE11 incision of the parental strand allows HR-mediated repair of the resulting structure (Figure 8B). Indeed, the 3’ end formed on the parental strand could invade the sister chromatid allowing repair from this template. Loss of HELQ may interfere with this process, since HELQ was shown to participate in D-loop extension as mentioned above (Adelman et al., 2013; Thomas et al., 2022; Ward et al., 2010). Thus, HELQ-depleted PRIMPOL-overexpressing cells not only accumulate ssDNA gaps due to defective gap filling, but also, upon MRE11-mediated conversion of these gaps into DSBs, are unable to repair these breaks since a critical step in the HR mechanism is compromised. DSB break accumulation eventually leads to cytotoxicity. Overall, our work suggests that as long as HR is intact, ssDNA gap accumulation may not necessarily cause cytotoxicity, which may potentially reconcile the findings that ssDNA gap accumulation correlates with chemosensitivity with the recent studies arguing that BRCA2 promotes therapy resistance primarily through HR (Feng and Jasin, 2017; Lim et al., 2024).

## Methods

### Cell culture and protein techniques

HeLa and U2OS cells (obtained from ATCC) were grown in Dulbecco’s modified Eagle’s media (DMEM). HeLa and U2OS PRIMPOL-overexpressing cells were generated in our laboratory and recently described (Nusawardhana et al., 2024). For PRIMPOL overexpression, the pLV[Exp]-Hygro-CMV>hPRIMPOL lentiviral construct (VectorBuilder) was used, while Empty-Vector (EV) control cells were obtained by infection with the pLV[Exp]-Hygro-CMV>ORF_Stuffer lentiviral construct (VectorBuilder). Infected cells were selected by hygromycin. HeLa-BRCA2^KO^ cells were generated in our laboratory and previously described (Clements et al., 2018). To knock-out HELQ, a commercially available CRISPR/Cas9 KO plasmid pRP[2CRISPR]-EGFP-hCas9-U6>hHELQ[gRNA]-U6>hHELQ[gRNA] (VectorBuilder) was used. Transfected cells were FACS-sorted into 96-well plates using a BD FACSAria II instrument. Resulting colonies were screened by Western blot.

Gene knockdown was performed using Lipofectamine RNAiMAX. AllStars Negative Control siRNA (Qiagen 1027281) was used as control. The following oligonucleotide sequences (Stealth or SilencerSelect siRNA, ThermoFisher) were used:

PRIMPOL: ID: 39536;
HELQ#1: CACAGAGAACCAGAGUGGAUAUGAA;
HELQ#2: ID:125860;
HELQ#3: ID: s41483;
NBN: ID: s529215;
PARP2: ID: s19504;
POLD3#1: ID: s21045;
POLD3#2: ID: 119736;
RAD52#1: ID: s11746;
RAD52#2: ID: s532174;
MUS81: UUUGCUGGGUCUCUAGGAUUGGUCU.

Denatured whole cell extracts were prepared by boiling cells in 100mM Tris, 4% SDS, 0.5M β-mercaptoethanol. Antibodies used for Western blot, at 1:500 dilution, were:

PRIMPOL (Proteintech 29824-1-AP);
HELQ: (Invitrogen PA5-88692);
POLD3: (Invitrogen PA5-96618);
RAD52: (Santa Cruz sc-365341);
MUS81: (Santa Cruz sc-47692);
BRCA2 (Bethyl A303-434A);
GAPDH (Santa Cruz Biotechnology sc-47724);
Vinculin (Santa Cruz Biotechnology sc-73614).

Inhibitors used were: RAD52 inhibitor -Epigallocatechin (EGC; Sigma-Aldrich, 970-74-1).

### CRISPR screens

For CRISPR knockout screens, the Brunello Human CRISPR knockout pooled lentiviral library (Addgene 73179) was used (Doench et al., 2016). This library encompasses 76,411 gRNAs that target 19,114 genes. 55 million cells from each cell lines (HeLa-EV, HeLa-PRIMPOL^OE^, U2OS-EV, U2OS-PRIMPOL^OE^) were infected with this library at a multiplicity of infection (MOI) of 0.4 to achieve 250-fold coverage and selected for 4 days with 0.6μg/mL puromycin. Twenty million library-infected cells (to maintain 250-fold coverage) were passaged for 14 days in the presence of 0.5μM cisplatin and then collected. Genomic DNA was isolated using the DNeasy Blood and Tissue Kit (Qiagen 69504) and employed for PCR using Illumina adapters to identify the gRNA representation in each sample. 10μg of gDNA was used in each PCR reaction along with 20μl 5X HiFi Reaction Buffer, 4μl of P5 primer, 4μl of P7 primer, 3μl of Radiant HiFi Ultra Polymerase (Stellar Scientific), and water. The P5 and P7 primers were determined using the user guide provided with the CRISPR libraries (https://media.addgene.org/cms/filer_public/61/16/611619f4-0926-4a07-b5c7-e286a8ecf7f5/broadgpp-sequencing-protocol.pdf). The PCR cycled as follows: 98°C for 2min before cycling, then 98°C for 10sec, 60°C for 15sec, and 72°C for 45sec, for 30 cycles, and finally 72°C for 5min. After PCR purification, the final product was Sanger sequenced to confirm that the guide region is present, followed by qPCR to determine the exact amount of PCR product present. The purified PCR product was then sequenced with Illumina HiSeq 2500 single read for 50 cycles, targeting 10 million reads. Next, the sequencing results were analyzed bioinformatically using the MAGeCK algorithm, which takes into consideration raw gRNA read counts to test if individual guides vary significantly between the conditions (Li et al., 2014). The MAGeCK software and instructions on running it were obtained from https://sourceforge.net/p/mageck/wiki/libraries/. Finally, analyses of the Gene Ontology and KEGG pathways enriched among the top hits was performed using DAVID (Ashburner et al., 2000; Huang da et al., 2009).

### Functional assays

Neutral and BrdU alkaline comet assays were performed (Thakar et al., 2020) using the Comet Assay Kit (Trevigen, 4250-050). For the BrdU alkaline comet assay, cells were incubated with 100μM BrdU as indicated. Chemical compounds (HU, cisplatin, olaparib) were added according to the labeling schemes presented. Slides were stained with anti-BrdU (BD 347580) antibodies and secondary AF568-conjugated antibodies (Invitrogen A-11031). Slides were imaged on a Nikon microscope operating the NIS Elements V1.10.00 software. Olive tail moment was analyzed using CometScore 2.0.

### Drug sensitivity assays

To assess cellular viability upon drug treatment, a luminescent ATP-based assay was performed using the CellTiterGlo reagent (Promega G7572) according to the manufacturer’s instructions. 1500 cells were seeded per well in 96-well plates and incubated as indicated. Luminescence was quantified using a Promega GloMax Navigator plate reader. For clonogenic survival assays, cells were treated with the indicated siRNA for 2 days, and 500 cells were seeded per well in 6-well plates and treated with the indicated dose of cisplatin. Media was changed after 7 days with fresh cisplatin at the indicated dose. After 10-14 days, colonies were washed with PBS, fixed with a solution of 10% methanol and 10% acetic acid, and stained with 2% crystal violet (Aqua solutions).

### DNA fiber combing assays

Cells were incubated with 100µM IdU and 100µM CldU as indicated. Chemical compounds (HU, cisplatin, olaparib) were added according to the labeling schemes presented. Next, cells were collected and processed using the FiberPrep kit (Genomic Vision EXT-001) according to the manufacturer’s instructions. Samples were added to combing reservoirs containing MES solution (2-(N-morpholino) ethanesulfonic acid) and DNA molecules were stretched onto coverslips (Genomic Vision COV-002-RUO) using the FiberComb Molecular Combing instrument (Genomic Vision MCS-001). For S1 nuclease assays, MES solution was supplemented with 1mM zinc acetate and either 40U/mL S1 nuclease (ThermoFisher 18001016) or S1 nuclease dilution buffer as control, and incubated for 30 minutes at room temperature. Slides were then stained with antibodies detecting CldU (Abcam 6236) and IdU (BD 347580), and incubated with secondary AF488 (Abcam 150117) or Cy5 (Abcam 6565) conjugated antibodies. Finally, the cells were mounted onto coverslips and imaged using a confocal microscope (Leica SP5) and analyzed using LASX 3.5.7.23225 software.

### Statistics and reproducibility

For DNA fiber assays and comet assays the Mann-Whitney statistical test (two-tailed) was performed. For CellTiterGlo cellular viability assays the two-way ANOVA statistical test was used. For clonogenic survival assays the t-test (two-tailed, unpaired) was used. For DNA fiber combing and comet assays, results from one experiment are shown; the results were reproduced in at least one additional independent biological conceptual replicate. Western blot experiments were reproduced at least two times. Statistical analyses were performed using GraphPad Prism 10 and Microsoft Excel v2205 software. Statistical significance is indicated for each graph (ns = not significant, for p>0.05; * for p≤0.05; ** for p≤0.01; *** for p≤0.001, **** for p≤0.0001). The random probabilities of identical genes within the top hits with MAGeCK score lower than 0.005 were calculated by multiplying the individual probabilities of each set: [(number of genes in set 1/total number of genes in the library) * (number of genes in set 2/total number of genes in the library)].

## Acknowledgements

We would like to thank Saiful Arefeen Sazed for technical support, and the Penn State College of Medicine Advanced Light Microscopy (RRID:SCR_022526), Genome Sciences (RRID:SCR_021123) and Flow Cytometry (RRID:SCR_021134) core facilities. Schematic figures were created with Biorender.com. This work was supported by: NIH R01ES026184 (to GLM), NIH R01GM134681 (to GLM), R01CA244417 (to CMN), NIH F31CA275340 (to LMP), as well as the Four Diamonds Transformative Patient-Oriented Cancer Research Project 4D01_2024_1002 (to GLM and CMN). The content is solely the responsibility of the authors and does not necessarily represent the official views of Four Diamonds.

## Author Contributions

L.M.P, C.M.N. and G.L.M. designed the experiments and wrote the paper. L.M.P and J.B.K. conducted the experiments.

