## Supplementary Figures for "CRISPR knockout genome-wide screens identify the HELQ-RAD52 axis in regulating the repair of cisplatin-induced single stranded DNA gaps"

#### Legends to Supplementary Tables

**Supplementary Table S1. MAGeCK analyses of the CRISPR screens identifying genes whose loss confers increased sensitivity to cisplatin in HeLa-EV cells.**

**Supplementary Table S2. MAGeCK analyses of the CRISPR screens identifying genes whose loss confers increased sensitivity to cisplatin in HeLa PRIMPOL-overexpressing cells.**

**Supplementary Table S3. MAGeCK analyses of the CRISPR screens identifying genes whose loss confers increased sensitivity to cisplatin in U2OS-EV cells.**

**Supplementary Table S4. MAGeCK analyses of the CRISPR screens identifying genes whose loss confers increased sensitivity to cisplatin in U2OS PRIMPOL-overexpressing cells.**

#### Legends to Supplementary Figures

**Supplementary Figure S1. Induction of ssDNA gaps in PRIMPOL-overexpressing cells.**

**A-D.** BrdU alkaline comet assays showing that PRIMPOL overexpression in HeLa (**A,C**) and U2OS (**B,D**) cells causes accumulation of replication-associated ssDNA gaps upon treatment with 0.4mM HU (**A,B**) or 10μM olaparib (**C,D**). At least 75 nuclei were quantified for each condition. The median values are marked on the graph and listed at the top. Asterisks indicate statistical significance (Mann-Whitney, two-tailed). Schematic representations of the assay conditions are shown at the top.

**E-G.** S1 nuclease DNA fiber combing assays showing that PRIMPOL overexpression in HeLa (**E,G**) and U2OS (**F,H**) cells causes accumulation of nascent strand ssDNA gaps upon treatment with 0.4mM HU (**E,F**) or 10μM olaparib (**G,H**). The ratio of CldU to IdU tract lengths is presented, with the median values marked on the graphs and listed at the top. At least 40 tracts were quantified for each sample. Asterisks indicate statistical significance (Mann-Whitney, two-tailed). Schematic representations of the assay conditions are shown at the top.

**Supplementary Figure S2. Analyses of CRISPR screens to identify genes causing cisplatin sensitivity to PRIMPOL-overexpressing cells.**

**A,B.** Biological pathway analyses using KEGG analyses of the top hits with MAGeCK score lower than 0.005 which cause cisplatin sensitivity in PRIMPOL-overexpressing HeLa (**A**) and U2OS (**B**) cells. KEGG terms with negative logP greater than 1.08 are presented.

**C.** Table showing the common hits (MAGeCK score lower than 0.005) in the HeLa and U2OS PRIMPOL-overexpressing cisplatin sensitivity screens, their ranks and their biological functions.

**Supplementary Figure S3. Analyses of CRISPR screens to identify genes causing cisplatin sensitivity to control (Empty Vector) cells.**

**A.** Diagram showing the overlap of identical genes within the top hits with MAGeCK score lower than 0.005 which cause cisplatin sensitivity in control (Empty Vector) HeLa and U2OS cells.

**B.** Table showing the common hits (MAGeCK score lower than 0.005) in the HeLa and U2OS control (Empty Vector) cisplatin sensitivity screens, their ranks and their biological functions.

**C.** The number of common genes within the top hits with MAGeCK score lower than 0.005 which cause cisplatin sensitivity in control (Empty Vector) HeLa and U2OS cells (namely 10) is slightly higher than the random probability of identical hits, which is 5.9.

**Supplementary Figure S4. Western blots showing siRNA-mediated knockdown of HELQ (A), POLD3 (B), RAD52 (C), and MUS81 (D) in HeLa cells.**

**Supplementary Figure S5. Impact of HELQ, RAD52 and POLD3 on ssDNA gap accumulation in PRIMPOL-overexpressing cells.**

**A.** BrdU alkaline comet assays showing that RAD52 depletion reduces ssDNA gap accumulation upon treatment with 10μM olaparib in PRIMPOL-overexpressing HeLa cells, while POLD3 depletion does not affect it. At least 110 nuclei were quantified for each condition. The median values are marked on the graph and listed at the top. Asterisks indicate statistical significance (Mann-Whitney, two-tailed). A schematic representation of the assay conditions is shown at the top.

**B,C.** S1 nuclease DNA fiber combing assays showing that RAD52 depletion suppresses ssDNA gap accumulation upon treatment with 150μM cisplatin in PRIMPOL-overexpressing HeLa (**B**) and U2OS (**C**) cells, while POLD3 depletion does not affect it. The ratio of CldU to IdU tract lengths is presented, with the median values marked on the graphs and listed at the top. The +S1/-S1 ratios of the median values are also presented. At least 85 tracts were quantified for each sample. Asterisks indicate statistical significance (Mann-Whitney, two-tailed). Schematic representations of the assay conditions are shown at the top.

**D.** S1 nuclease DNA fiber combing assays showing that HELQ depletion does not impact ssDNA gap accumulation upon treatment with 150μM cisplatin in PRIMPOL-overexpressing BRCA2-knockout HeLa cells. The ratio of CldU to IdU tract lengths is presented, with the median values marked on the graphs and listed at the top. The +S1/-S1 ratios of the median values are also presented. At least 100 tracts were quantified for each sample. Asterisks indicate statistical significance (Mann-Whitney, two-tailed). A schematic representation of the assay conditions is shown at the top.

### Supplementary Figure S1

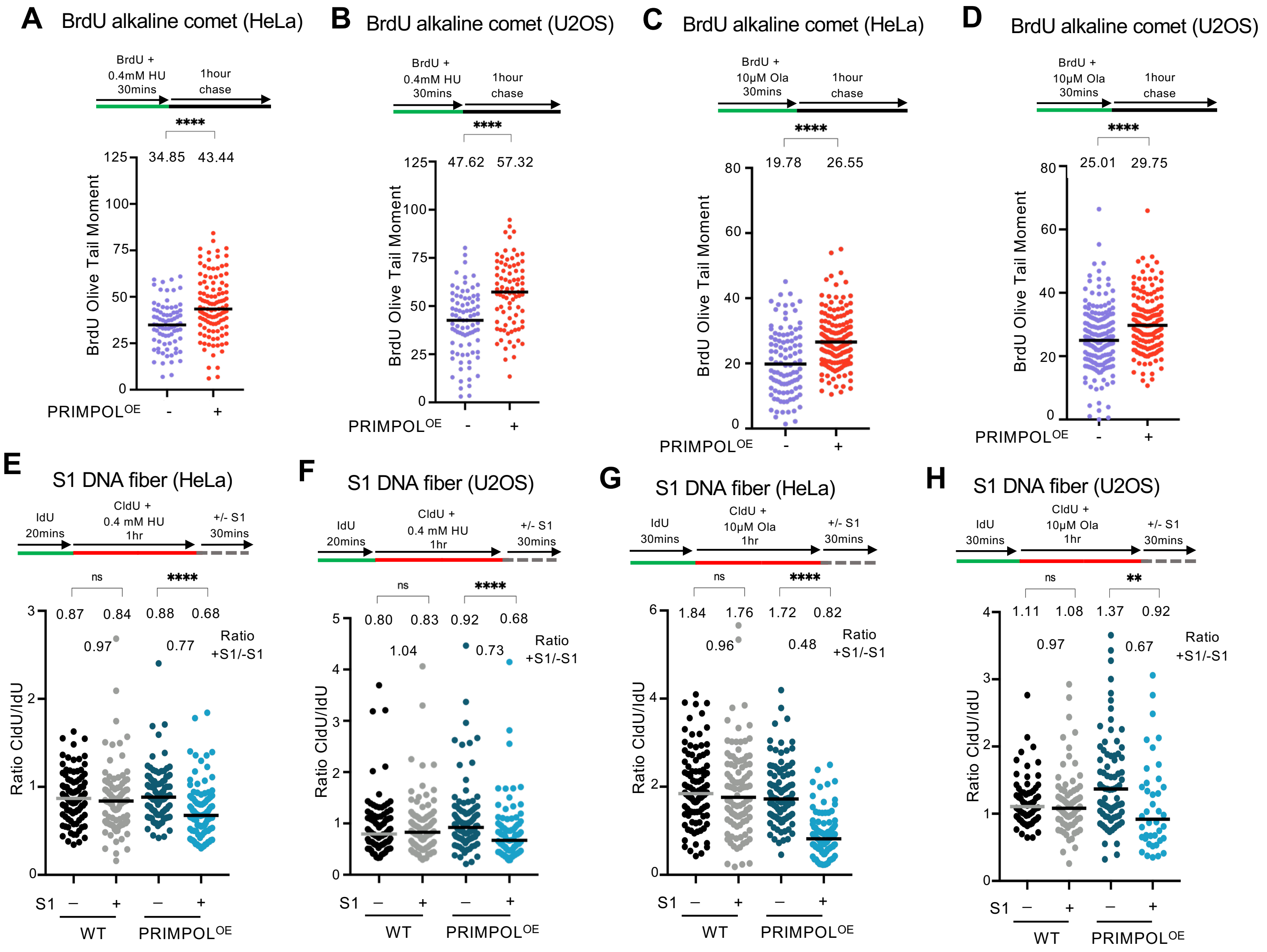

### Supplementary Figure S2

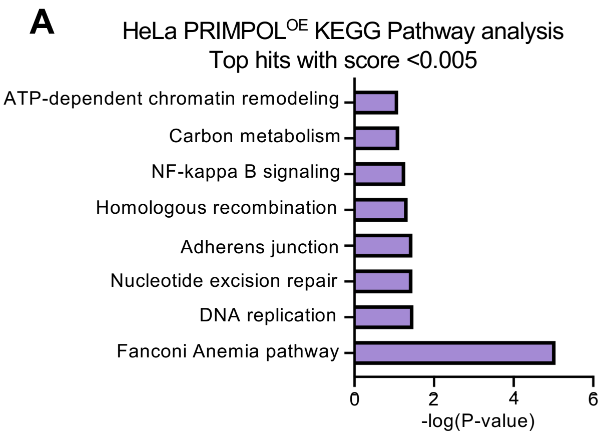


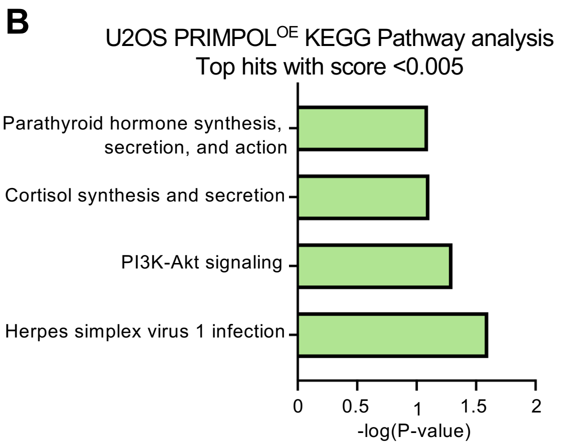


**C**

| Gene | Rank Cisplatin sensitivity HeLa PRIMPOL$^{OE}$ | Rank Cisplatin sensitivity U2OS PRIMPOL$^{OE}$ | Function |
|---|---|---|---|
| HELQ | 174 | 191 | ssDNA ATPase and DNA helicase |
| GDI1 | 166 | 6 | GDP dissociation inhibitor in GDP-GTP exchange reaction |
| THNSL2 | 211 | 16 | Threonine synthase-like protein with phospho-lyase activity |
| OR10W1 | 140 | 68 | Olfactory receptor protein |
| GPN1 | 304 | 132 | Guanosine triphosphatase enzyme that may play a role in DNA repair |
| STAB1 | 188 | 169 | Transmembrane receptor with potential functions in angiogenesis, lymphocyte homing, cell adhesion |
| PRDX2 | 111 | 212 | Antioxidant enzyme |
| ZFP62 | 169 | 246 | Zinc finger protein with DNA-binding transcription activator activity |
| SLC35E2 | 159 | 257 | Solute carrier predicted to be involved in transmembrane transport |
| OR8J3 | 285 | 279 | Olfactory receptor protein |
| DIRAS2 | 84 | 324 | Ras monomeric GTPase |

#### Supplementary Figure S3

**A**

Empty Vector cisplatin sensitivity screens
Common genes among the top hits with score <0.005

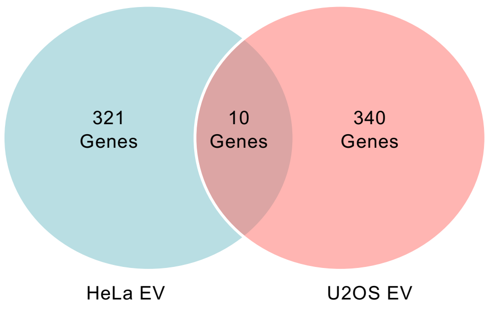


**C**

PRIMPOL$^{OE}$ cisplatin sensitivity screens
Random probability of identical genes among
top hits with score <0.005

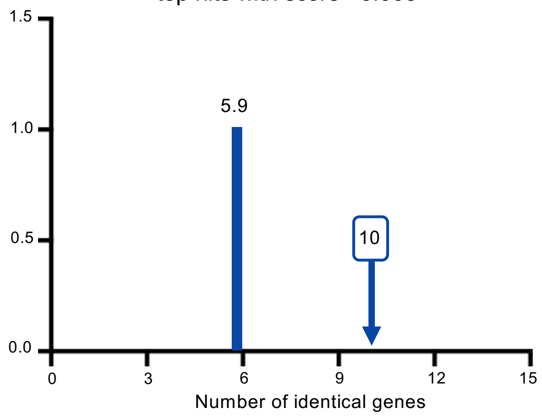


**B**

| Gene | Rank Cisplatin sensitivity HeLa EV | Rank Cisplatin sensitivity U2OS EV | Function |
|---|---|---|---|
| ATM | 51 | 164 | DNA damage checkpoint activation |
| PAPPA2 | 90 | 334 | Metzincin metalloproteinase; cleaves insulin growth factor-binding protein 5 |
| BEST2 | 98 | 159 | Bestrophin family anion channel |
| SMTNL1 | 109 | 21 | Striated and smooth muscle contraction |
| TMEM255B | 116 | 332 | Integral membrane component |
| RTN4RL1 | 168 | 303 | Potential negative regulation of axon regeneration |
| DTX3L | 215 | 170 | E3 Ubiquitin ligase; STAT protein binding activity |
| CASC3 | 250 | 158 | Component of exon junction complex; nonsense-mediated mRNA decay |
| HAS2 | 329 | 77 | Hyaluranic acid synthase |
| FANCA | 91 | 320 | DNA repair; Interstrand DNA cross-link repair |

#### Supplementary Figure S4

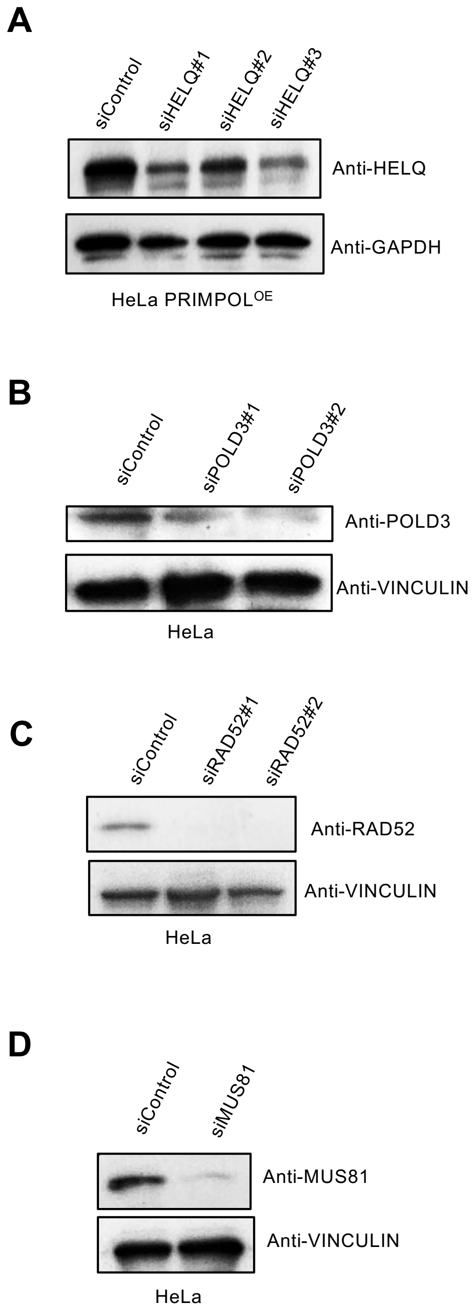

### Supplementary Figure S5

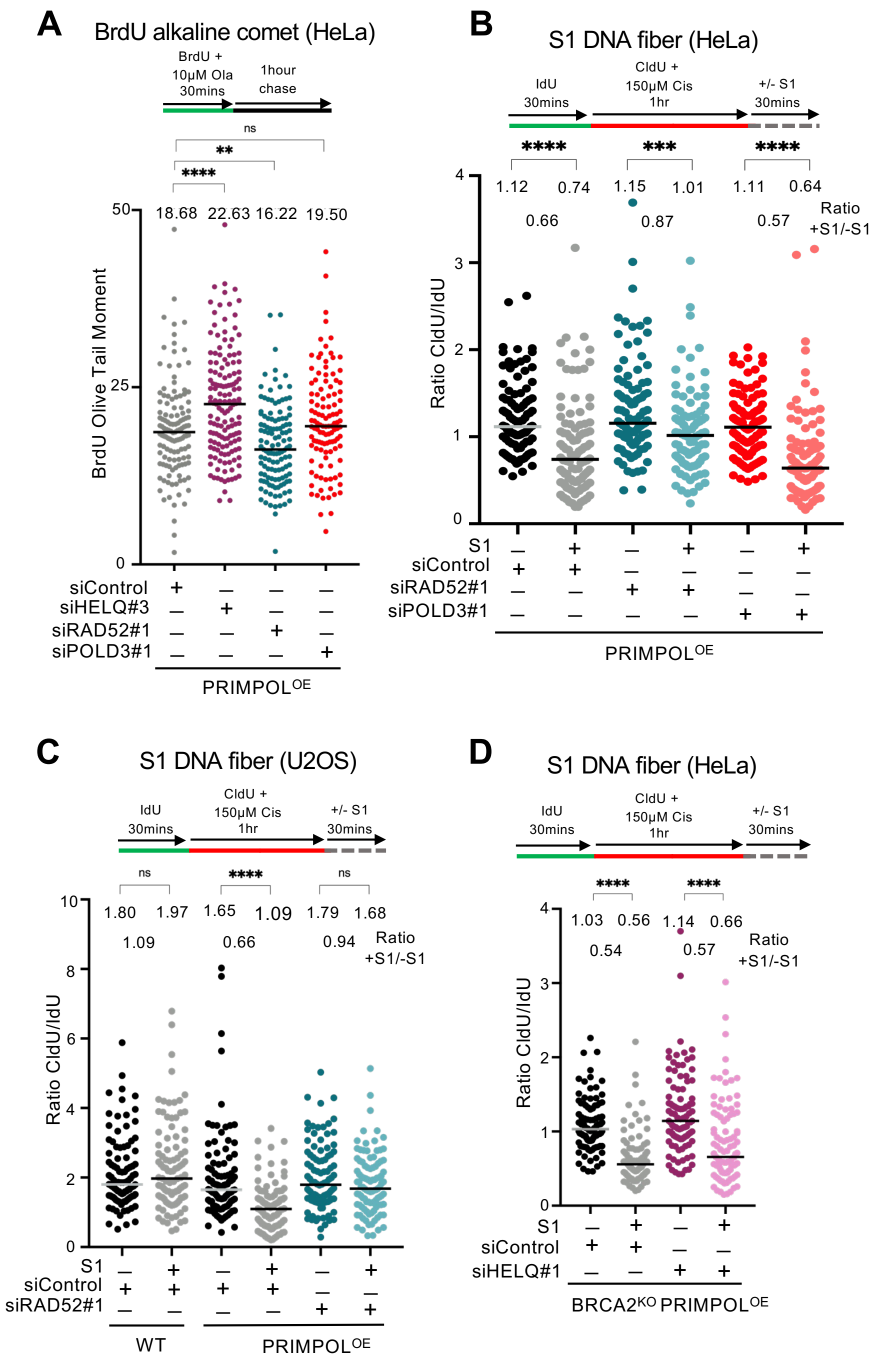
